# Pre-existent adult sox10^+^ cardiomyocytes contribute to myocardial regeneration in the zebrafish

**DOI:** 10.1101/662536

**Authors:** Marcos Sande-Melón, Inês J. Marques, María Galardi-Castilla, Xavier Langa, María Pérez-López, Marius Botos, Gabriela Guzmán-Martínez, David Miguel Ferreira-Francisco, Dinko Pavlinic, Vladimir Benes, Remy Bruggmann, Nadia Mercader

## Abstract

During heart regeneration in the zebrafish, fibrotic tissue is replaced by newly formed cardiomyocytes derived from pre-existing ones. It is unclear whether the heart is comprised of several cardiomyocyte populations bearing different capacity to replace lost myocardium. Here, using *sox10* genetic fate mapping, we identified a subset of pre-existent cardiomyocytes in the adult zebrafish heart with a distinct gene expression profile that expanded massively after cryoinjury. Genetic ablation of *sox10*^+^ cardiomyocytes severely impaired cardiac regeneration revealing that they play a crucial role for heart regeneration.

## Introduction

Cardiomyocyte (CM) renewal in the human heart is marginal and, after acute myocardial infarction, millions of CMs are irreversibly lost and replaced by a fibrotic scar (Prabhu and Frangogiannis, 2016). Adult mammalian CMs have a poor capacity to proliferate and an efficient contribution of an adult stem cell pool for myocardial replacement has not been demonstrated (Lerman, et al., 2016). By contrast, zebrafish have an extraordinary capacity for heart regeneration after injury (Gonzalez-Rosa, et al., 2017; Kikuchi, 2014). Lineage tracing studies have revealed that pre-existing CMs are the origin of *de novo* formed cardiac muscle (Jopling, et al., 2010; Kikuchi, et al., 2010). Upon injury, CMs adjacent to the injury lose their sarcomeric organization and exhibit a more immature phenotype. Concomitant with this structural change, CMs begin to express developmental genes, suggesting a reversion of their differentiated phenotype to a more embryonic-like state (Lepilina, et al., 2006). Indeed, *gata4*, *hand2* and *tbx5a*, which play key roles during heart development (Grajevskaja, et al., 2018; Garrity, et al., 2002; Kuo, et al., 1997; Molkentin, et al., 1997; Srivastava, et al., 1997), are required for heart regeneration (Grajevskaja, et al., 2018; Schindler, et al., 2014; Gupta, et al., 2013). Moreover, CMs contributing to heart regeneration activate *gata4* and *ctgfa* enhancer elements upon injury (Gupta, et al., 2013; Kikuchi, et al., 2010)(Pfefferli and Jazwinska, 2017). While CMs that will contribute to heart regeneration upregulate a specific set of genes, it is unclear whether CM subsets contributing to regeneration can be distinguished by means of its expression profile in the uninjured heart.

Like mammals, zebrafish CMs derive from first and second heart field progenitors (Mosimann, et al., 2015; Zhou, et al., 2011; de Pater, et al., 2009). However, in the zebrafish, the neural crest represents a third progenitor population that contributes to the developing heart. Cell transplantation and fluorescent dye tracing experiments suggested that cranial neural crest cells incorporate not only into the areas of the outflow tract, as in mammals and birds, but also into the atrium and ventricle (Li, et al., 2003; Sato and Yost, 2003). Moreover, genetic lineage tracing using *sox10* as a neural crest cell marker revealed a cellular contribution of *sox10*^+^ cells to the zebrafish heart (Cavanaugh, et al., 2015; Mongera, et al., 2013) and suggested that *sox10*-derived CMs are necessary for correct heart development (Abdul-Wajid, et al., 2018). Noteworthy, it is still unclear if a *sox10*^+^ neural crests population differentiates into CMs or if a *sox10*^+^ CM subset is relevant for heart development.

Here we assessed the contribution of *sox10*-derived cells to the adult zebrafish heart using *sox10*:CreER^T2^ fate mapping during homeostasis and regeneration. We found that embryonic *sox10*-derived cells contributed to significant portions of the adult heart. We also identified adult sox10^+^ CMs that expanded to a higher degree upon injury than the rest of CMs and which significantly contributed to cardiac regeneration. Their transcriptome differed from other CMs in the heart and their genetic ablation impaired the recovery from ventricular cryoinjury.

### Embryonic *sox10*^+^ cells contribute to the adult heart

*sox10*-lineage tracing has revealed the presence of *sox10*-derived CMs in the developing zebrafish heart (Abdul-Wajid, et al., 2018; Cavanaugh, et al., 2015; Mongera, et al., 2013). Recently, the presence of adult *sox10*-derived cells has been reported in the adult zebrafish heart (Abdul-Wajid, et al., 2018). However, the lack of an inducible lineage tracing system and therefore non-temporary control of recombination generates doubts about the developmental origin of these cells in the adult heart. To clarify the source of *sox10-*derived cells contributing to the adult heart, we crossed Tg(*sox10:CreER^T2^*) (Mongera, et al., 2013) with Tg(*ubb:loxP-GFP-loxP-mCherry)* (Mosimann, et al., 2011), from now on named *ubb:Switch*. In double transgenic animals, 4-Hydroxytamoxifen (4-OHT) administration leads to constitutive m^i^Cherry expression in cells expressing *sox10:CreER^T2^* at the time of recombination. The *sox10:CreER^T2^* line additionally harbors a myocardial reporter cassette: a *myosin light chain 7* (*myl7*) promoter element driving GFP expression in CMs. Embryos were treated with 4-OHT between 12 and 48 hours post-fertilization (hpf) (Fig. 1A), the time window spanning neural crest cell migration and heart tube formation. Live imaging was performed at 5 days post-fertilization (dpf). We detected mCherry^+^/*myl7*:GFP^+^ in the larval heart, indicating the presence of a subset of CMs derived from *sox10*-expressing cells (Fig. 1B,C n=7). After live imaging, the larvae were grown separately to adulthood to follow the fate of mCherry^+^ cells. In the adult, we again detected mCherry^+^ cell clusters, showing that embryonic *sox10*-derived cells contribute to the adult heart (Fig. 1D-F n=7). We observed that these cells contributed to 5% of the total ventricular volume (Fig. 1F-H). Most of the mCherry^+^ cells were also *my7l*:GFP^+^ and thus CMs (Fig. 1I). Immunostaining with the myocardial marker Myosin Heavy Chain (MHC) confirmed that the *sox10*-derived cells were CMs (Fig. 1J,K). As reported for *sox10:Cre* genetic tracing (Abdul-Wajid, et al., 2018), we detected more *sox10*-derived cells in the trabecular myocardium than in the compact myocardium using *sox10:CreER^T2^* (Fig. 1L).We quantified the distribution of *sox10*-derived CMs in the adult hear. We observed a predominant contribution to the basal and medial myocardium-regions close to the atrioventricular canal (AVC)-as well as to a lesser extent to the apical myocardium (most distant to the AVC) (Fig. 1M; n=19 hearts analyzed).

**Figure 1.**
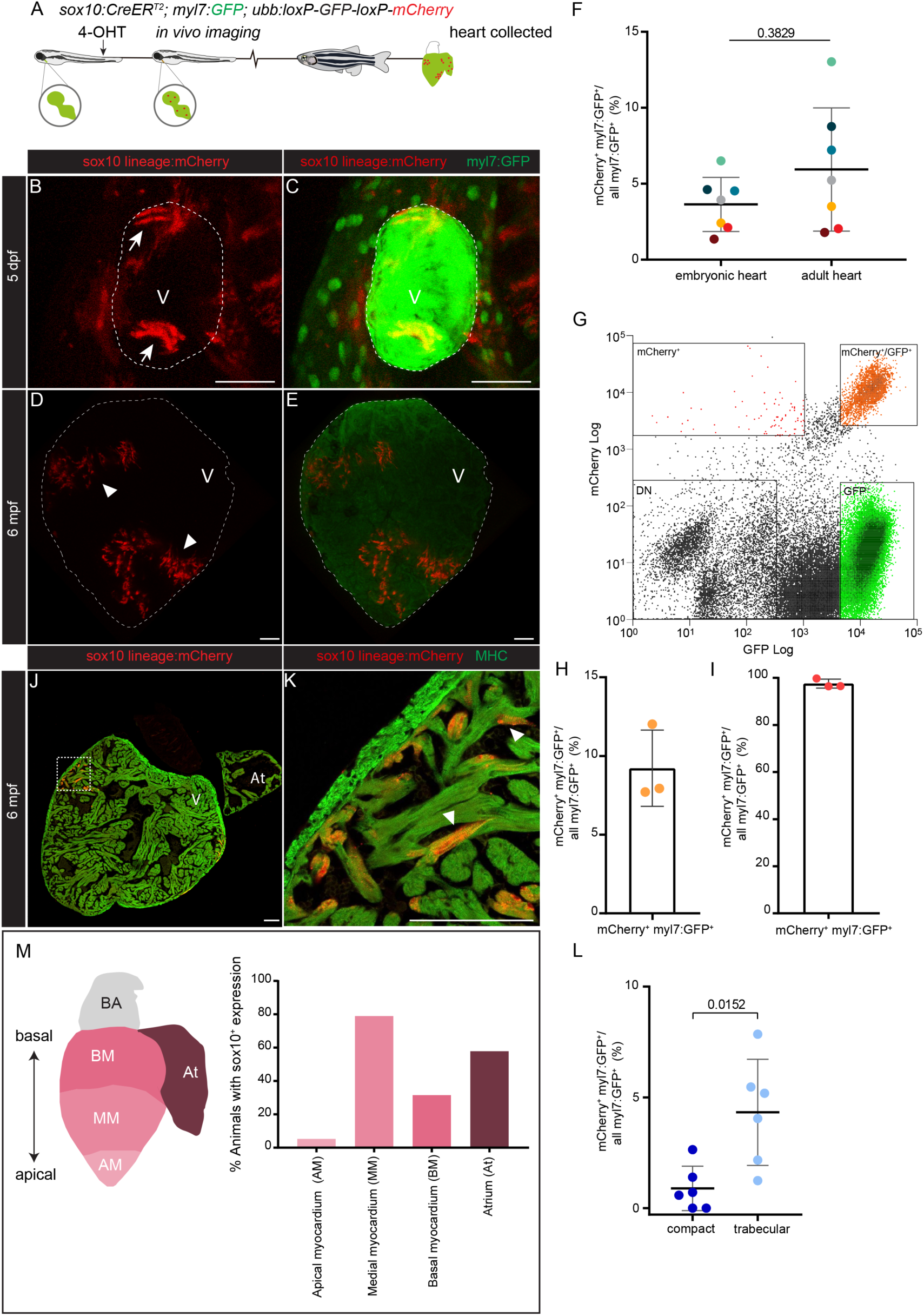
Embryonic *sox10*-derived cells contribute to the adult myocardium. (A), *sox10:CreER^T2^;ubb:loxP-GFP-loxP-mCherry* embryos were treated with 4-OHT from 12 to 48 hours post-fertilization. *sox10*-derived cells are labelled with mCherry. These cardiac cells were detected at 5 dpf by confocal live imaging. Larvae were grown to adulthood and hearts collected to characterize the contribution of embryonic mCherry^+^ cells to the adult heart. (B-C) *sox10*-derived cells within the embryonic heart. Maximal projection of a confocal z-stack through the heart at 5 days post-fertilization (dpf); shown is a ventral view of the ventricle. Anterior is to the top. Cardiomyocytes (CMs) are shown in green, *sox10*-derived CMs in red. (D-E) Embryonic *sox10*-derived cells contribute to the adult heart. Maximal projection of a confocal z-stack through the adult ventricle corresponding to embryonic heart shown in (B,C). (F) Quantification of the percentage of mCherry^+^/MHC^+^ from the total MHC^+^ volume at 5 dpf and adult ventricles. Colored dots mark individual animals (n=7). Data are means ± s.d.; p=0.3289 (two-tailed non-parametric t-test). (G-H) Representative example of a FAC sorting experiment of embryonic *sox10*-derived cells from adult hearts. Gates show *sox10*-derived CMs as orange dots (*myl7*:GFP^+^ mCherry^+^), *myl7*:GFP^+^ single positive cells as green dots (GFP^+^) and the rest of *sox10*-derived cells (mCherry^+^) as red dots. 4 different hearts were pooled. (H) Percentage of FAC-sorted *sox10*-derived CMs (mCherry^+^, *my7*:GFP^+^) vs the total amount of myocardium (*myl7*:GFP^+^). (I) Percentage of mCherry^+^ *myl7*:GFP^+^ volume *vs* the total number of mCherry^+^ cells in the adult heart. (J) Immunostaining of a sagittal heart section for MHC (green) and mCherry (*sox10* lineage; red). (K) Zoomed view of boxed area in (K). (L) Percentage of *sox10*-derived CMs in the compact and trabecular myocardium of embryonic hearts at 5 dpf quantified on whole mount images. Shown are data from individual hearts as well as means ± s.d.; p= 0.0152 (two-tailed non-parametric t-test). (M) Localization of embryonic *sox10*-derived CMs in the adult heart (n=19). Percentage of fish, which after embryonic recombination show mCherry^+^ CMs in the adult heart. The cardiac ventricle is outlined by dotted lines in (B-E). Basal Myocardium category (BM include the atrioventricular canal myocardium). Arrows point to *sox10*-derived CMs clusters. 4-OHT, 4-hydroxytamoxifen; dpf, days post-fertilization; MHC; myosin heavy chain; mpf, months post-fertilization; V, ventricle. Scale bars: 100 µm.

To determine whether the embryonic *sox10*-derived population expanded in response to injury, we cryoinjured ventricles from adult Tg(*sox10:CreER^T2^*);(*ubb:Switch*) zebrafish recombined during embryogenesis and compared the percentage of mCherry^+^ cells in injured and uninjured hearts (Fig. S1A). mCherry^+^ cells were detected in a similar proportion in uninjured (Fig. S1B-E, n=12) and injured hearts at 14 days post-injury (dpi) (Fig. S1F-J, n=9). Analysis of the proportion of mCherry^+^ CMs contributing to different regions from the injury site showed that there was no preferred contribution to any particular region (Fig. S1K; n= 5). Similarly, in uninjured hearts, no apicobasal region was identified with a preferred contribution of mCherry^+^ cells (Fig. S1L; n= 5). These results imply that in the adult heart, CMs derived from *sox10*^+^ embryonic cells do not expand in response to injury and are not preferentially contributing to myocardium regeneration.

### A subset of adult *sox10*-expressing cells contributes extensively to the regenerated myocardium

We next wanted to assess whether there were *sox10*^+^ cells also in the adult heart (Fig. 2A). Administration of 4-OHT to adult *Tg*(*sox10:CreER^T2^*);(*ubb:Switch*) zebrafish led to the activation of mCherry expression in a few cardiac cells, scattered throughout the ventricle (Fig. 2B-E; n= 6). Whole-mount immunofluorescence confirmed the presence of *sox10*-derived CMs (Fig. B-B’’’). These were present in the cortical as well as in the peripheral trabecular layer (Fig. 2D-E).

**Figure 2.**
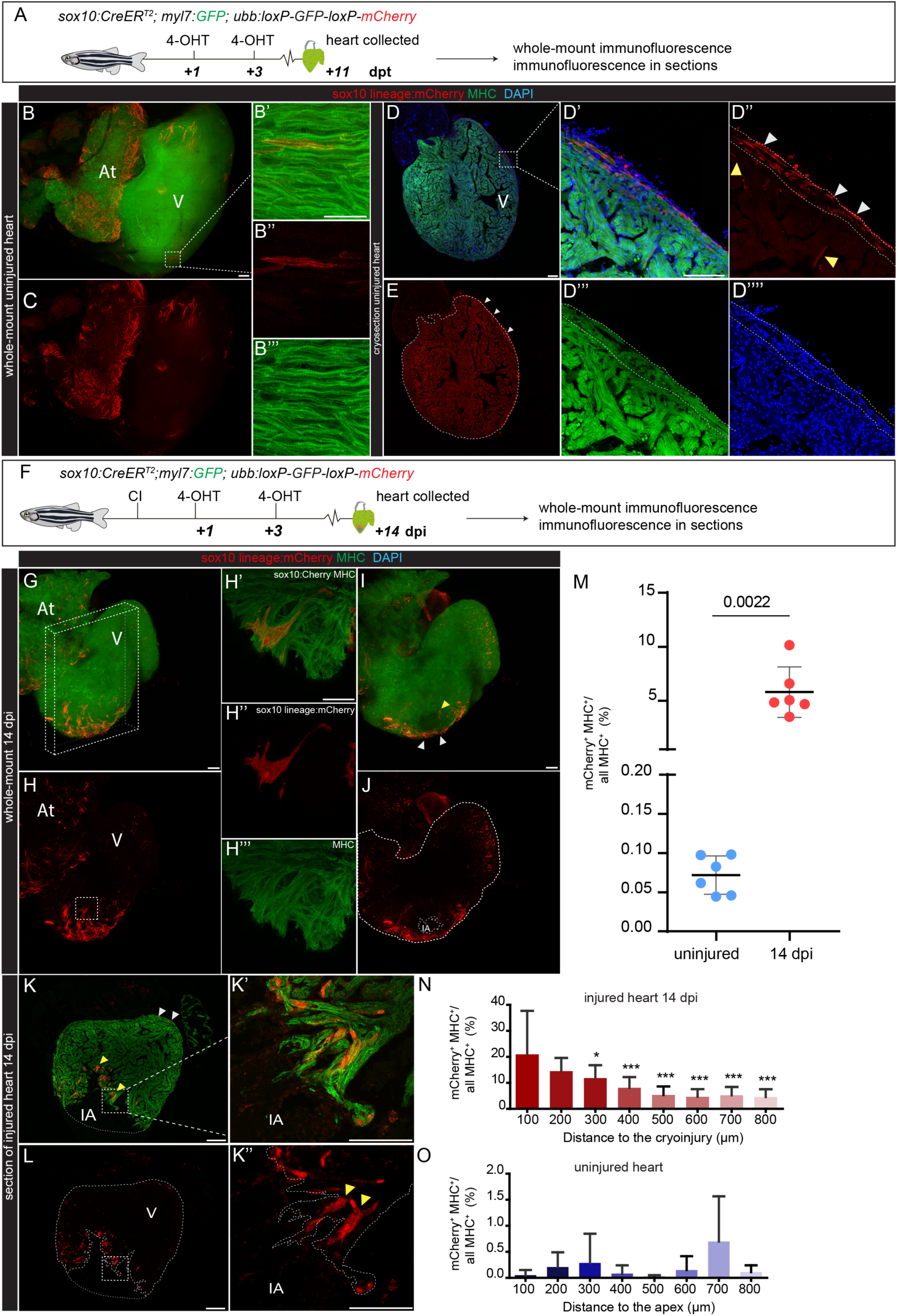
Contribution of *sox10*-derived CMs to heart regeneration. (A) Two 4-OHT pulses were administered to adult *sox10:CreER^T2^;ubb:loxP-GFP-loxP-mCherry* zebrafish 11 and 9 days before heart extraction to induce mCherry in *sox10:CreER^T2^* expressing cells. mCherry (red), *sox10* lineage. anti-MHC, marks CMs (green) in uninjured hearts (B-E) or hearts at different days postinjury (G-L).- (B-C) Whole-mount view of uninjured hearts (Maximal projection of 352 µm, 44 z-planes). (B’-B’’’) Maximum projection of zoomed view of boxed area in B (7 µm, 7 z-planes). (D-E) Immunostaining on cryosection of an uninjured heart. (D’-D’’’’) Zoomed view of boxed area in (D). *sox10*-derived CMs and non-CMs are observed in the subepicardial area of the ventricle. mCherry^+^ cell in the cortical myocardium (white arrowheads), mCherry^+^ trabecular CMs (yellow arrowheads). (F) Two 4-OHT pulses were administered to adult *sox10:CreER^T2^;ubb:loxP-GFP-loxP-mCherry* one and three days after the injury to induce mCherry expression in *sox10:CreER^T2^* expressing cells. Hearts were extracted at 14 dpi. (G-J) Whole-mount view of injured hearts at 14 dpi (594 µm, z-73 planes). *sox10*-derived CMs can be observed in the apical region of the heart. (H’-H’’’) Zoomed view of boxed area in (H). (I-J) z-stack from heart in G (252 µm, 31 z-planes), showing injured area. Yellow arrowheads, mCherry^+^; MHC^+^ trabecular CMs. White arrowheads, mCherry^+^; MHC^+^ cortical CMs. (K-L) Paraffin sections of injured hearts at 14 dpi. Note the distribution of *sox10*-derived CMs around the IA (yellow arrowheads). (K-K’) Paraffin section of injured heart at 14 dpi and zoomed view of cortical (white arrowheads) and trabecular (yellow arrowheads) *sox10*-derived CMs adjacent to the IA. (M) Percentage of the volume from mCherry^+^/MHC^+^ cells relative to all MHC^+^ cells in uninjured (n=6) and injured hearts (n=6) at 14 dpi. The volume of *sox10*-derived CMs expands after injury. Data are means ± s.d.; p= 0.0022 (two-tailed non-parametric t-test). (N) The ventricles from a whole-mount stained hearts were digitally sectioned in increments of 100 µm starting from the injury site. Shown is the percentage of the volume from mCherry^+^/MHC^+^ cells relative to all MHC^+^ cells within different heart segments. The percentage of mCherry^+^/MHC^+^ is high near to the injury site and decreases towards ventricular regions that are distant (injured hearts n=6). Data are means ± s.d.; *p<0.05; ***p<0.001 (one-way ANOVA test). (O) The ventricle was digitally sectioned in increments of 100 µm starting from the apex. Shown is the percentage of the volume from mCherry^+^/MHC^+^ cells relative to all MHC^+^ cells within different heart segments. The percentage of mCherry^+^/MHC^+^ is similarly distributed through ventricular regions of the uninjured ventricle heart (uninjured hearts n=6). Data are means ± s.d.; p>0.05 (one-way ANOVA test). 4-OHT, 4-hydroxytamoxifen; At, atrium; CI, cryoinjury; CMs, cardiomyocytes; dpi, days post-injury; dpt, days post-treatment; IA, injured area; V, ventricle. Scale bars: 100 µm.

The presence of adult *sox10*^+^ CMs was confirmed using the line Tg*(−4.9sox10:GFP*)*^ba2^* (Carney, et al., 2006). We were able to detect few *sox10*:GFP^+^ CMs in the ventricle of injured and uninjured hearts (Figure S2A-E’’; n= 4 hearts). Moreover, RNAScope mRNA detection revealed *sox10* expression in few cells close to the cortical and trabecular myocardial boundaries, as well as at the borders of the injury area (Fig. S2 F-I’’; n=4 uninjured, n=4 injured).

Since embryonic *sox10*-derived cells did not seem to contribute to heart regeneration, we analyzed if this was also the case for the adult *sox10*^+^ cells. When Tg(*sox10:CreER^T2^*);(*ubb:Switch*) zebrafish were treated with 4-OHT shortly after ventricular cryoinjury (Fig. 2F), we indeed observed an expansion of *sox10*-derived cells (Fig. 2G-L; n= 6). Immunostaining on whole hearts and on sections confirmed that *sox10*-derived cells were CMs, as they co-expressed MHC (Fig. 2H’-2H’’’ and Fig. 2K’-K’’). In order to estimate the contribution of *sox10*-derived CMs during the regeneration process, we measured the ratio of *sox10*-derived CMs (mCherry^+^/MHC^+^) versus the rest of CMs (MHC^+^) in the entire heart (Fig. 2M). When we compared the proportion of mCherry^+^ ventricular CMs from cryoinjured and control hearts, we observed that the portion of mCherry^+^ myocardium increased >100-fold at 7dpi (Fig. 2M). We then questioned whether *sox10*-derived cells expanded globally in the injured heart or whether there was a distinct contribution to the regenerating myocardium. To do this, we generated a distribution map of distance for the *sox10*-derived CMs from the injury site (for injured hearts, Fig. 2N) or apical myocardium (for uninjured hearts, Fig. 2O) to the basal myocardium. The closer to the injury area the greater was the contribution of *sox10*-derived CMs (Fig. 2N, n= 6). This bias towards an apical region was not detected in uninjured hearts (Fig. 2O; n= 6).

Surprisingly, we did not detect an expansion of *sox10*:GFP^+^ cells after cardiac injury (Figure S2A-E’’; n=4 hearts). This result might indicate that the *sox10*-derived cells in regenerating hearts were descendants from a small subset of sox10^+^ CMs within the uninjured heart that turned off *sox10* expression after expansion.

### Pre-existent *sox10*-derived CMs contribute to cardiac regeneration

To further assess whether the increased number of *sox10*-derived CMs is a result of the expansion of a small pool of pre-existent *sox10*^+^ CMs, we induced recombination in adult zebrafish before cryoinjury (Fig. 3 and Fig S2J-T). We administered 4-OHT to adult zebrafish, either few days (Fig S2) or two weeks before the injury (Fig. 3A). 4-OHT treatments at 12 and 14 days prior to cardiac injury allowed to rule out that non-metabolized 4-OHT could be active and induce recombination after injury. We detected *sox10*-derived cells in the atrium, ventricle and valves of uninjured hearts (Fig. 3B-D’’). We detected both, *sox10*-derived CMs (mCherry^+^/MHC^+^) as well as non-CMs (only mCherry^+^) (Fig. 3C’,C’’ and 3D’,D’’). At 7 dpi, *sox10*-derived CMs were present mainly close to the injury and in subepicardial areas both, in the trabecular and cortical myocardium (Fig. 3E). Indeed, both in injured and uninjured heart, *sox10*-derived cells were preferentially detected in the compact myocardium (Fig. S2P). In both regions, the proportion of *sox10*-derived CMs significantly expanded upon injury (Fig. S2P and 3F-G). Interestingly, the percentage of *sox10*-derived CM is higher close to injury area compared to the basal ventricle (Fig. 3G). This preferential distribution of *sox10*-derived CMs towards the apical region could not be observed in uninjured hearts of siblings (Fig. 3G).

**Figure 3.**
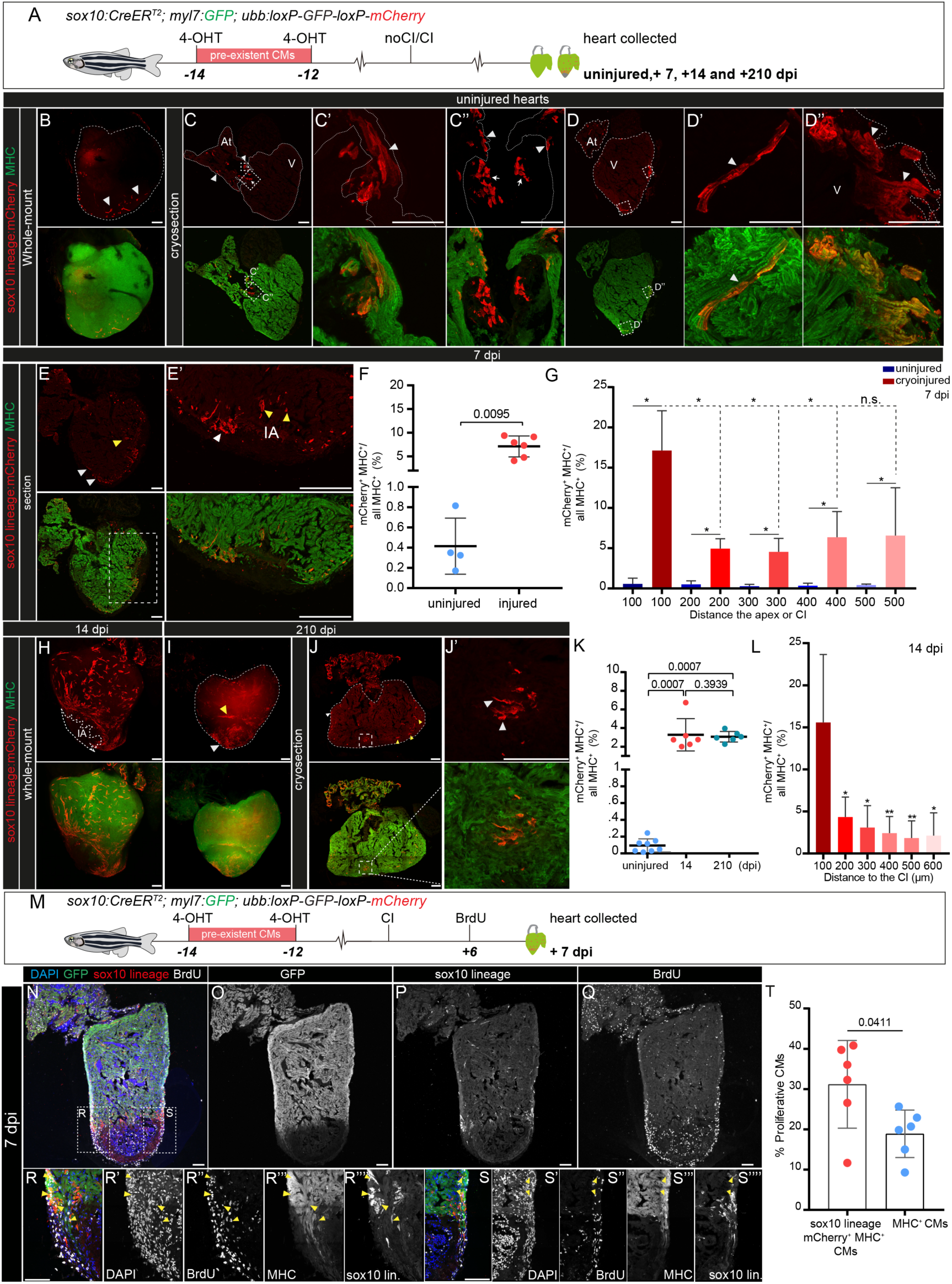
Pre-existent *sox10*-derived CMs expand at the injury area and contribute to cardiac regeneration. (A) Adult *sox10:CreER^T2^;ubb:loxP-GFP-loxP-mCherry* zebrafish were treated with 4-OHT on day 14 and 12 before cryoinjury. Hearts were fixed at 7, 14 or 210 dpi and processed for immunostaining with anti-mCherry^+^ (red, *sox10*-derived cells) and anti-MHC (green, myocardium). (B-D’’) Whole-mounts (B) and cryosections (C-D’’) of uninjured hearts. Upper row, mCherry channel, lower row, merged channels. Arrowheads in C, *sox10*-derived cells. C’-D”, zoomed views of panels C and D. Arrowheads, mCherry^+^ CMs; arrows, mCherry^+^ non-CMs in within the atrioventricular valve. D-D’’ mCherry^+^ CMs in the medial (D’’) and apical regions (D’) of the ventricle ventricle. (E-E’) Cryosection of an injured heart at 7 dpi. *sox10*-derived CMs are detected near the IA and subepicardial regions of the myocardium. White arrowheads, mCherry^+^ CMs in the cortical myocardium, yellow arrowheads, mCherry^+^ CMs in the trabecular myocardium. (F) Quantification of the proportion of *sox10*-derived CMs in uninjured (n=4) and injured hearts (n=6) at 7 dpi. Each dot represents the value from one heart. Shown are also means ± s.d.; p= 0.0095 (two-tailed non-parametric t-test). (G) Quantification of the distribution of ventricular mCherry^+^ CMs on uninjured and cryosectioned hearts at 7 dpi. The ventricle was digitally divided into increments of 100 µm starting from the injury site in injured hearts or apex in uninjured hearts. Shown is the percentage of the mCherry^+^/MHC^+^ area relative to the whole MHC^+^ area within different heart segments. The percentage of mCherry^+^/MHC^+^ is high near to the IA and decreases towards ventricular regions distant to the IA (injured hearts n=5; uninjured hearts, n=4). Statistical differences represented in dashed lines, injured hearts represented in red bars. The percentage of mCherry^+^/MHC^+^ is higher in every region of the ventricular heart of injured hearts (red bars) compared to uninjured hearts (blue bars). Data are means ± s.d.; *p<0.05 (two-tailed non-parametric t-test). (H) Whole-mount image of a heart at 14 dpi. *sox10*-derived CMs are distributed around the IA and distant part of the ventricle. (I-J’) Whole-mount view (I) and cryosections (J-J’) of regenerated hearts at 210 dpi. *sox10*-derived CMs can be observed in the apical region of the heart. White arrowheads, mCherry^+^ CMs in the cortical myocardium, yellow arrowheads, mCherry^+^ CMs in the trabecular myocardium. (K) Quantification of the percentage of mCherry^+^ CMs in uninjured cardiac ventricles compared to ventricles at 14 and 210 dpi (whole mount immunostained hearts). Shown are measurements of individual hearts (dots) as well as means ± s.d. p=0.3939 p= 0.007 (two-tailed non-parametric t-test; uninjured n= 8, 14 dpi n=6, 210 dpi n= 6). (L) Quantification of the distribution of mCherry^+^ CMs in whole mount immunostained hearts. Distance calculation as shown in H. Data are means ± s.d. *p<0.05 (two-tailed non-parametric t-test; n= 6). (M-O) Assessment of the proliferation index of *sox10*-derived CMs. Two pulses of 4-OHT were administered, 14 and 12 days before the injury. BrdU was added at 6 dpi and hearts collected at 7 dpi. (N-Q) Heart section immunostained with anti-BrDU (white), anti-mCherry (red, *sox10* lineage) and MHC (green). Nuclei were counterstained with DAPI (blue). (R-S’’’) Zoomed views of boxed regions in N. Yellow arrowheads, mCherry^+^,BrdU^+^, MHC^+^ triple positive cells. (T) Quantification of mCherry^+^,BrdU^+^, MHC^+^ triple positive cells versus all BrdU^+^,MHC^+^ double positive cells in the 100 µm IA border zone. Shown are values for individual hearts as well as means ± s.d. *p<0.05 (two-tailed non-parametric t-test; n= 6). At, Atrium, dpi, days postinjury; IA, injured area; MHC, myosin heavy chain; V, Ventricle. Scale bars: 100 µm except for 4E’ (200 µm).

We also analyzed the proportion of *sox10*-derived cells at later stages of regeneration to assess if the expansion is temporal or whether these cells are contributing to the fully regenerated heart (Fig. 3H-L). At 14 dpi, *sox10*-derived CMs remained around the injured area (Fig. 3H). Even at 210 dpi, when the regeneration is complete(Gonzalez-Rosa, et al., 2011), mCherry^+^/MHC^+^ CMs were still detected within the region presumably corresponding to the regenerated myocardium (Fig 3J-J’). Quantification showed that the mCherry^+^/MHC^+^ myocardial volume was significantly higher in injured than in uninjured hearts at all regeneration stages analyzed (Fig. 3K). Consistent with the results at 7dpi, the proportion of the mCherry^+^ myocardial volume was significantly higher in areas adjacent to the injury than in peripheral regions at 14 dpi (Fig. 3L).

To better understand the mechanisms of the accumulation of *sox10*-derived cells at the injury area, we assessed cell proliferation (Fig. 3M-T). Recombination was induced at 14 and 12 days before cryoinjury and BrdU injected at 6 dpi (Fig. 3M). Immunostaining against mCherry, to detect the *sox10* lineage, the myocardial marker MHC, and BrdU to label proliferating cells, showed a statistically significant increase in the amount of proliferating *sox10*-derived CMs (n=6) when compared to the remaining CMs (n=6) (Fig. 3T). Thus, CMs with an active *sox10* promoter element in the uninjured heart divide at a higher rate than the rest of CMs in response to injury.

When performing the experiment in juveniles, we did observe a qualitative increase in *sox10*-derived cell area after injury (Fig S3A-T). However, quantification of the signal on sections did not yield a significant difference (Fig. S3T). Importantly, control experiments, in which no 4-OHT was added, yielded no recombination and the heart were completely devoid of mCherry expression, both in juveniles and adults (Fig. S3U-W). Thus, with sox10:CreER^T2^ we are fully controlling recombination events and can therefore faithfully trace the fate of cells with active *sox10*:*CreER* ^T2^ expression at the time of 4-OHT addition.

Collectively, these results show that the *sox10:CreER^T2^;ubb:Switch* double transgenic line enables the detection of a subset of pre-existent CMs that expands in response to injury and contributes preferentially to myocardial regeneration in the zebrafish.

### *sox10*-derived cardiomyocytes reveal a specific gene signature

To investigate if *sox10*-derived CMs differ in their gene expression profile compared to the rest of CMs, we performed a transcriptome analysis (Fig. 4A,B). To make the characterization specific for the myocardium, we used the line *sox10:CreER^T2^*;*vmhcl:loxP-tagBFP-loxP-mCherry-NTR,* in which upon 4-OHT induced recombination, *sox10*-derived CMs express mCherry and the rest of ventricular CMs blue fluorescent protein (BFP). Two pulses of 4-OHT, 14 and 12 days before the injury, were administered. Hearts of uninjured and regenerating *sox10:CreER^T2^*;*vmhcl:loxP-tagBFP-loxP-mCherry-NTR* animals were collected, disgregated and mCherry^+^, BFP^+^ as well as double positive mCherry^+^/BFP^+^ CMs were FAC sorted. Samples consisting of 20 CMs each were processed for RNA sequencing (RNA-Seq). For bioinformatics analysis, we compared mCherry^+^ (comprised of all samples that were mCherry^+^ or mCherry^+^/BFP^+^) with mCherry^-^ (comprised of samples that were only BFP^+^). We observed that in uninjured hearts, 101 genes were upregulated and 129 genes were downregulated in sox10-derived (mCherry^+^) CMs compared to the rest of ventricular CMs (mCherry^−^) (Fig. 4A, B Fig S4).

**Figure 4.**
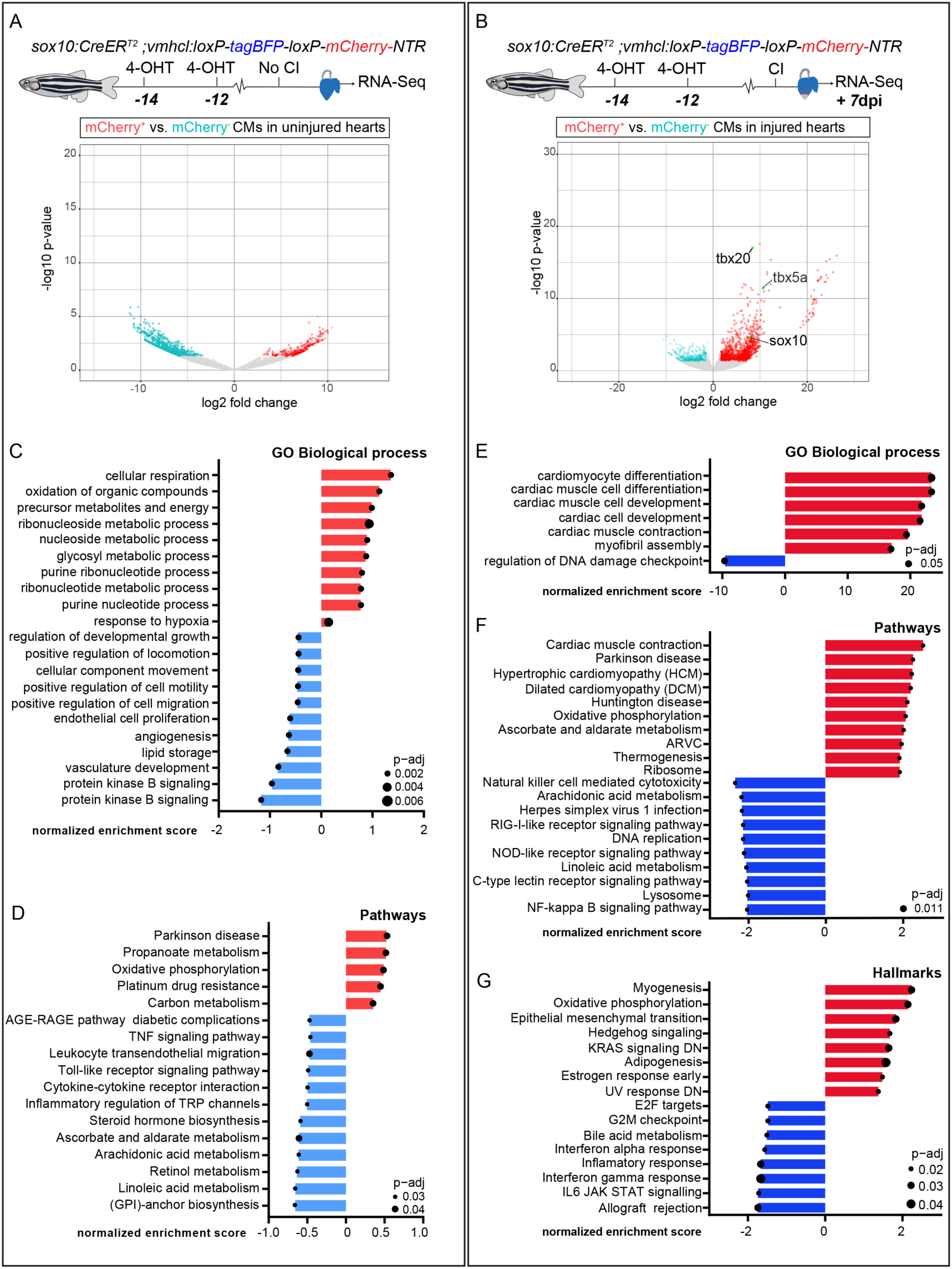
*sox10*-derived CMs reveal a specific transcriptome signature. Hearts from *sox10:CreER^T2^*;*vmhcl:loxP-tagBFP-loxP-mCherry-NTR* zebrafish treated with 4-OHT were collected 12 days after 4-OHT treatment. Hearts were disgregated, *sox10*-derived CMs (mCherry^+^) and rest of CMs of the heart (mCherry^-^) were FAC-sorted, and RNA-Seq was performed on 26 samples consisting of 20 CMs each. Recombination and collection was carried out either on uninjured hearts (A) or hearts at 7dpi (B). (A) Volcano plots representing differentially expressed genes (DEG) in Cherry^+^ and Cherry^-^ CMs from uninjured (A) or 7 dpi (B) hearts. DEG upregulated in Cherry^-^ CMs are blue, and DEG upregulated in Cherry^+^ CMs are red. These genes had an adjp-value ≤0.05 and a log2 fold change (LFC) ≥ 2. Genes with an FDR ≥0.05 and LFC ≤ ± 2 are represented in grey. Note the high number of DEG upregulated in mCherry^+^ cells at 7dpi. Some differentially expressed genes previously linked to regeneration as well as *sox10* are highlighted. (C) Biological processes differentially enriched in Cherry^+^ vs Cherry^-^ CMs from uninjured hearts (source for analysis DEG list). (D) Pathway analysis using Gene enrichment analysis (GSEA) of genes with LFC ≥ 2 present in Cherry^+^ and Cherry^-^ CMs of uninjured hearts. Representative biological processes were plotted and ordered according to the normalized enrichment score. Z-score represents whether a specific function is increased or decreased. Red bar, enriched in mCherry^+^ CMs, blue bar, enriched in rest of CMs. (E) Biological processes differentially enriched in Cherry^+^ vs Cherry^-^ CMs from uninjured hearts (source for analysis DEG list). (F,G) Pathway analysis and Hallmarks using Gene enrichment analysis (GSEA) of genes with LFC ≥ 2 present in Cherry^+^ and Cherry^-^ CMs of injured hearts. Representative biological processes were plotted and ordered according to the normalized enrichment score. Z-score represents whether a specific function is increased or decreased. Red bar, enriched in mCherry^+^ CMs, blue bar, enriched in rest of CMs. 4-OHT, 4-Hydroxytamoxifen; CI, cryoinjury; dpi, days post-injury; dpt, days post-treatment; FDR, false discovery rate; GO, Gene Ontology.

Gene enrichment analysis of mCherry^+^ and mCherry^-^ transcriptomic profiles in uninjured hearts revealed several metabolic differences between these two groups including changes in oxidative phosphorylation and nucleic acid metabolism (Fig. 4C-D).

Notably, after injury, when comparing mCherry^+^ with mCherry^-^ groups we found that *sox10*-derived cells were transcriptionally more active: 415 genes were upregulated in mCherry^+^ CMs while only 30 genes were upregulated in mCherry^-^ CMs. Importantly, *sox10* mRNA was significantly upregulated in mCherry^+^ CMs from injured hearts at 7 dpi, showing that the *sox10:CreER^T2^* line can be used to trace endogenous *sox10* expressing cells (Fig. 4B). The genes encoding the T-box transcription factors *tbx20* as well as *tbx5a* were among the genes upregulated in *sox10*-derived CMs (Fig. 4B). Theses genes were previously shown to be expressed in CM populations involved in heart regeneration and to play an active role in this process (Grajevskaja, et al., 2018; Xiang, et al., 2016).

Gene enrichment analysis in injured conditions for mCherry^+^ and mCherry^-^ CMs showed that Gene Ontology (GO) Biological Processes related to negative regulation of the cell cycle were inhibited in mCherry^+^ CMs. This suggests that mCherry^+^ CMs have a pro-regenerative profile (Fig. 4E-G). Furthermore, mCherry^+^ CMs were enriched for pathways involved in myocardial growth including CM differentiation, cardiac cell development, or cardiac muscle contraction (Fig. 4E-G). This result is consistent with a role for *sox10*-derived CMs in rebuilding the lost myocardium.

We next analysed if mCherry^+^ cells respond equally to an injury than the rest of CMs. For this, we compared on the one hand the changes in gene expression of mCherry^+^ CMs in injured with uninjured hearts (Fig. S5) and on the other the changes in gene expression of mCherry^-^ CMs between both conditions (Fig. S6). Upon injury, mCherry^+^ cells upregulated 767 genes and only 26 genes were downregulated (Fig. S5A,B). Thus overall, mCherry^+^ CMs respond to injury with an increase in gene expression. Biological processes related to cell proliferation, cell motility, and response to injury were enriched in mCherry^+^ CMs upon cryoinjury (Fig. S5C). Ingenuity pathway (IPA) further confirmed the enrichment of canonical pathways related to proliferation in mCherry^+^ CMs of injured hearts (Fig. S5D).

When we analysed the changes in gene expression of the rest of CMs (mCherry^-^) in response to injury, we observed fewer DEG than those detected for mCherry^+^ cells (Fig S6). This indicated that *sox10*-derived cells reactivate gene expression to a larger extent compared to the rest of CMs.

Altogether, this data indicates that *sox10*-derived CMs not only preferentially contribute to heart regeneration, but that *sox10* promoter expression also defines a group of myocardial cells in the adult uninjured zebrafish heart with a unique gene expression signature and pro-regenerative transcriptomic profile in response to injury.

### *sox10*^+^ cells are necessary for cardiac regeneration

The accumulation of *sox10*-derived cells in the regenerated myocardium strongly suggests that these cells contribute to the replacement of injured myocardium. To determine the function of this population during heart regeneration, we genetically ablated *sox10*-derived cells using the transgenic line *sox10:CreER^T2^*;*β-actin:loxPmCherryloxP-DTA* (Wang, et al., 2011) by administration of 4-OHT to adults 3 and 1 days before cryoinjury (Figure S7A). At 21 dpi, the injured area was larger in *sox10*^+^ cell-depleted animals than in the control group (Figure S7B–D). There was no significant reduction in animal survival between both groups (Figure S7E) and cardiac function was equal to animals from the control group (Figure S7 F-I; n=6). During development, *sox10*-derived cells contribute to the peripheral nervous system, which plays an important role in heart regeneration (Mahmoud, et al., 2015; White, et al., 2015). Thus, the impaired regeneration in animals with *sox10*-ablated cells might be a consequence of compromised cardiac innervation. Yet, the *β-actin* promoter used in the transgenic line has been reported to be strongly expressed in CMs but weak in the rest of cardiac cells (Kikuchi, et al., 2010). Furthermore, we did not observe a reduction in innervation in cryoinjured hearts when compared to controls (Fig. S7J-N).

To test the effect of ablation of *sox10*-derived myocardial cells more specifically, we genetically ablated *sox10*-derived cells in ventricular CMs using *sox10:CreER^T2^*;*vmhcl:loxP-tagBFP-loxP-mCherry-NTR* (Sanchez-Iranzo, et al., 2018) zebrafish (Fig. 6A). With this double transgenic line ventricular *sox10*^+^-derived CMs can be genetically ablated upon addition of Metronidazole (Mtz), which induces cell death in nitroreductase (NTR)-expressing cells (Curado, et al., 2008) (Fig. 5A). We ablated mCherry^+^ cells one week before cryinjury and confirmed efficiency of ablation by comparing the amount of mCherry^+^ cells with Mtz non-treated animals (Fig. 5B,C). A significant difference in the amount of *sox10*-derived CMs could be observed between untreated (n=5) and treated (n=8) adult zebrafish (Fig. 5D), confirming the efficiency of ablation and a lack of repopulation of mCherry^+^ cells during regeneration. Consistent with the results using the genetic ablation model by DTA, the survival rate of the different groups was not statistically significant (Fig. 5E). To assess if loss of adult *sox10*-derived ventricular CMs affects heart regeneration after cryoinjury, we collected hearts at 30 dpi and assessed fibrotic tissue deposition (Fig. 5 F-H). Histological staining on heart sections revealed a persistent fibrotic scar and a larger injury area in fish in which ventricular *sox10*-derived CMs had been ablated (Fig. 5F), when compared to the control groups *sox10:CreER^T2^*;*vmhcl:loxP-mCherry-NTR* without Mtz administration (Fig. 5G) and *sox10:CreER^T2^*;*ubb:loxP-mCherry* with Mtz (Fig. 5H) (Fig. 5I). This indicates that ablation of the small pool of *sox10*-derived ventricular CMs in the uninjured adult zebrafish heart, comprising less than 1% of total myocardial volume, affects subsequent heart regeneration.

**Figure 5.**
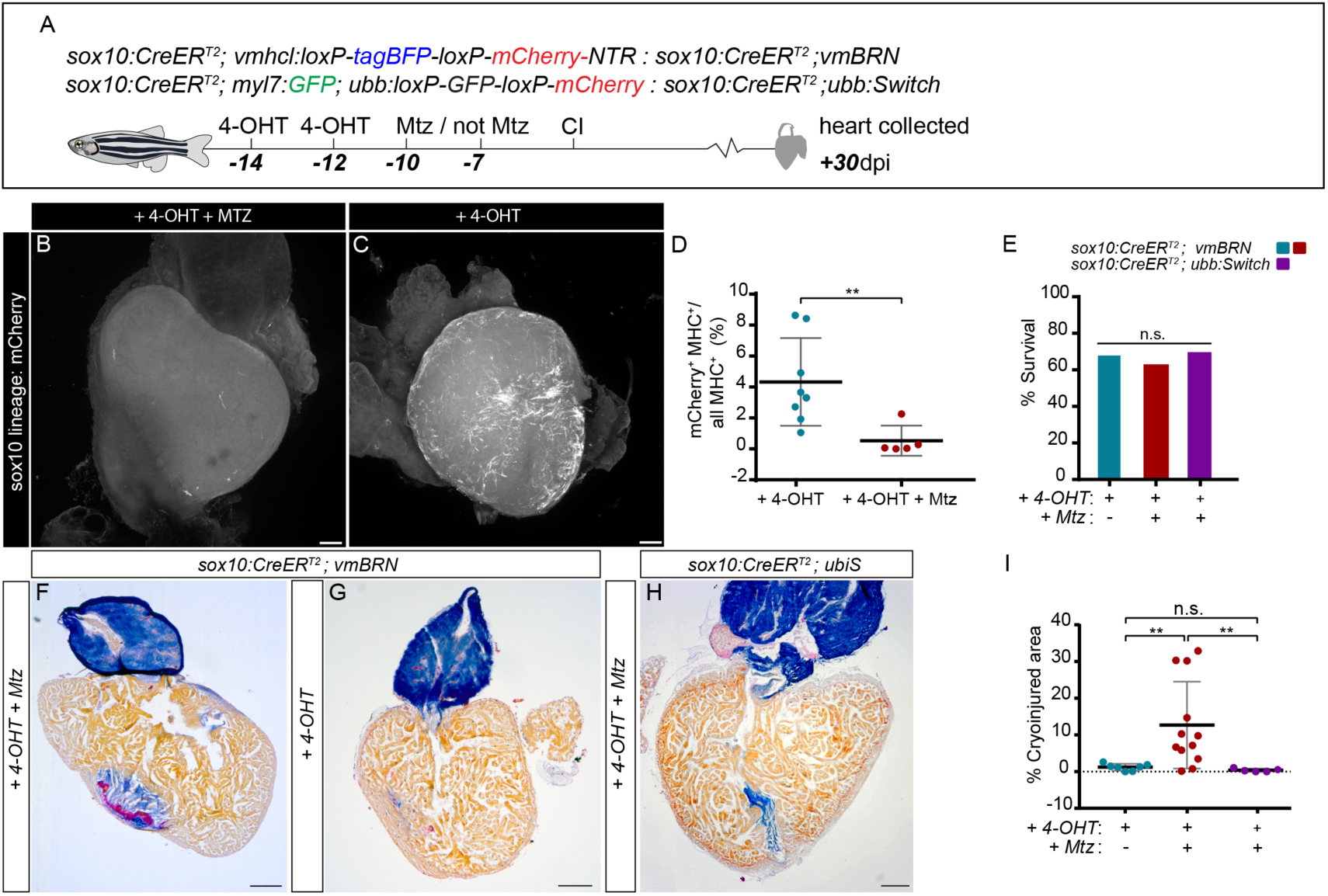
Genetic ablation of *sox10*^+^ cardiomyocytes impairs cardiac regeneration. (A) At days −14 and −12 prior to cryoinjury, *sox10:CreER^T2^*;*vmBRN* or *sox10:CreER^T2^*;*ubb:Switch* zebrafish were treated with 4-OHT. They were treated with Mtz on days 10 and 7 before injury. A control group of *sox10:CreER^T2^*;*vmBRN* was not treated with Mtz. Cryoinjured hearts were collected at 30 dpi. (B,C) Whole mount view of confocal 3D projection of z-stacks through a *sox10:CreER^T2^*;*vmBRN* heart after 4-OHT and with (B) or without (C) Mtz treatments and 30 dpi. No mCherry^+^ cells are observed in (B), revealing efficient ablation of *sox10*-derived cells. Recombined cardiomyocytes (CMs) are visible close to the site of regeneration in (C). (D) Percentage of the volume from mCherry^+^ CMs relative to all CMs: mCherry^+^; Myosin Heavy Chain (MHC)^+^ *vs* all MHC^+^ cells (p=0.006; two-tailed non-parametric t-test). (E) Survival rate of animals from groups as described in A. No difference in mortality was observed among the groups according to a Fisher’s exact test (p=1.000). (F-H) AFOG histological staining on sagittal sections of cryoinjured hearts at 30 dpi, (F) 4-OHT and Mtz-treated *sox10:CreER^T2^*;*vmBRN* heart section. (G) 4-OHT-treated *sox10:CreER^T2^*;*vmBRN* heart section. (H) 4-OHT and Mtz-treated *sox10:CreER^T2^*;*ubb:Switch* heart section. (I) Quantification of IA in the three conditions shown in E-F. IA versus total ventricular myocardial area was measured. Shown are values for individual hearts as well as means ± s.d. Statistical analysis: non-parametric t-test. *sox10:CreER^T2^*;*vmBRN* Mtz-treated *vs. sox10:CreER^T2^*;*vmBRN Mtz-untreated* (p= 0.0078); *sox10:CreER^T2^*;*vmBRN Mtz-treated vs. sox10:CreER^T2^*;*ubb:Switch* Mtz-treated (p=0.0072); *sox10:CreER^T2^*;*vmBRN* Mtz-untreated vs. *sox10:CreER^T2^*;*ubb:Switch* Mtz-treated (p=0.1020). 4-OHT, 4-hydroxytamoxifen; At, atrium; CI, cryoinjury; dpi, days post-injury; IA, injured area; Mtz, Metronidazole V, ventricle. Scale bars: B,C 100 µm; F-H, 200 µm.

Collectively, these data strongly suggest that ventricular *sox10*-derived CMs in the adult zebrafish heart are required for cardiac regeneration.

## Discussion

CMs can proliferate in the adult zebrafish (Wills, et al., 2008), and this is likely the basis for the high regenerative capacity observed in the injured heart. Although little is known about CM populations that contribute to regeneration, studies showed that they activate *gata4* and *ctgfa* regulatory elements in response to injury (Kikuchi, et al., 2010) (Pfefferli and Jazwinska, 2017). Recent clonal analysis studies using pan-myocardial lineage tracing have suggested that distinct CM subset can contribute to heart regeneration both in the zebrafish and in the mouse (Sereti, et al., 2018; Gupta, et al., 2013). Here we report for the first time genetic fate mapping with a specific promotor element of pre-existent CMs, which are present in the adult heart and expand more than the rest of CMs in response to injury. Our study suggests that this small *sox10*-derived CM population is essential for regeneration, since its genetic ablation prior to cryoinjury impairs cardiac regeneration.

Adult neural crest stem cells have been recently proposed as the source of progenitor cells during adult pigment cell regeneration in the zebrafish (Iyengar, et al., 2015). Moreover, in rodents, neural crest stem cells have been suggested to participate in repair mechanisms after myocardial infarction (Tamura, et al., 2016). The results presented here do not fully support a neural crest cell-origin of the *sox10*^+^ population contributing to heart regeneration as we did not observe contribution to the regenerating heart of embryonic *sox10*^+^ cells. Instead, our results suggest the presence of a small pool of *sox10*^+^ CMs in the adult heart that efficiently expands in response to injury. Adult *sox10*^+^ CMs reveal a unique gene signature, both in uninjured hearts and upon injury. Interestingly, *sox10* transcripts were also detected in CMs in a recently published single-cell transcriptome of zebrafish embryos (Wagner, et al., 2018), further supporting our findings of a *sox10*^+^ CM population in the zebrafish heart. Moreover, here we also report the expression of endogenous *sox10* expression in *sox10*-derived CMs, supporting that our driver lines recapitulate expression of the endogenous gene. The biological pathways enriched in *sox10*-derived CMs compared to the rest of ventricular myocardium were related to developmental processes, metabolism and cell proliferation. This gene signature could be key for their increased contribution to the regenerated myocardium. The fact that *sox10*-derived cells express more genes in response to injury that the rest of ventricular CMs could suggest that they are epigenetically less repressed and therefore more sensitive to injury response and prone to contribute to heart regeneration.

CMs with active *sox10* promoter expression might represent a particular state of CMs or mark a distinct CM population in the zebrafish heart with high regenerative capacity. Understanding if this population is unique to this species or shared in mammals might help to understand the basis of regenerative capacity.

### STAR Methods

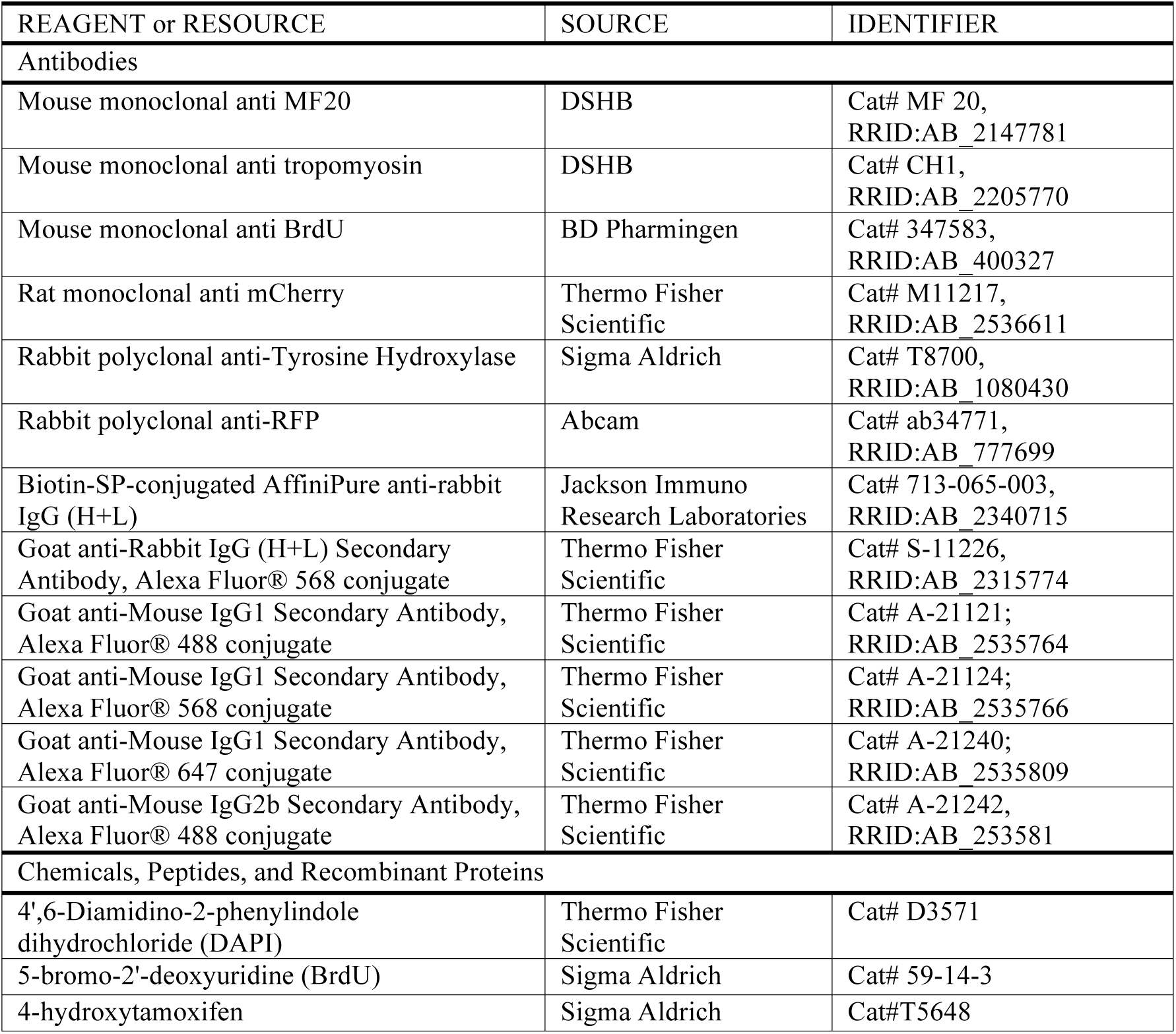
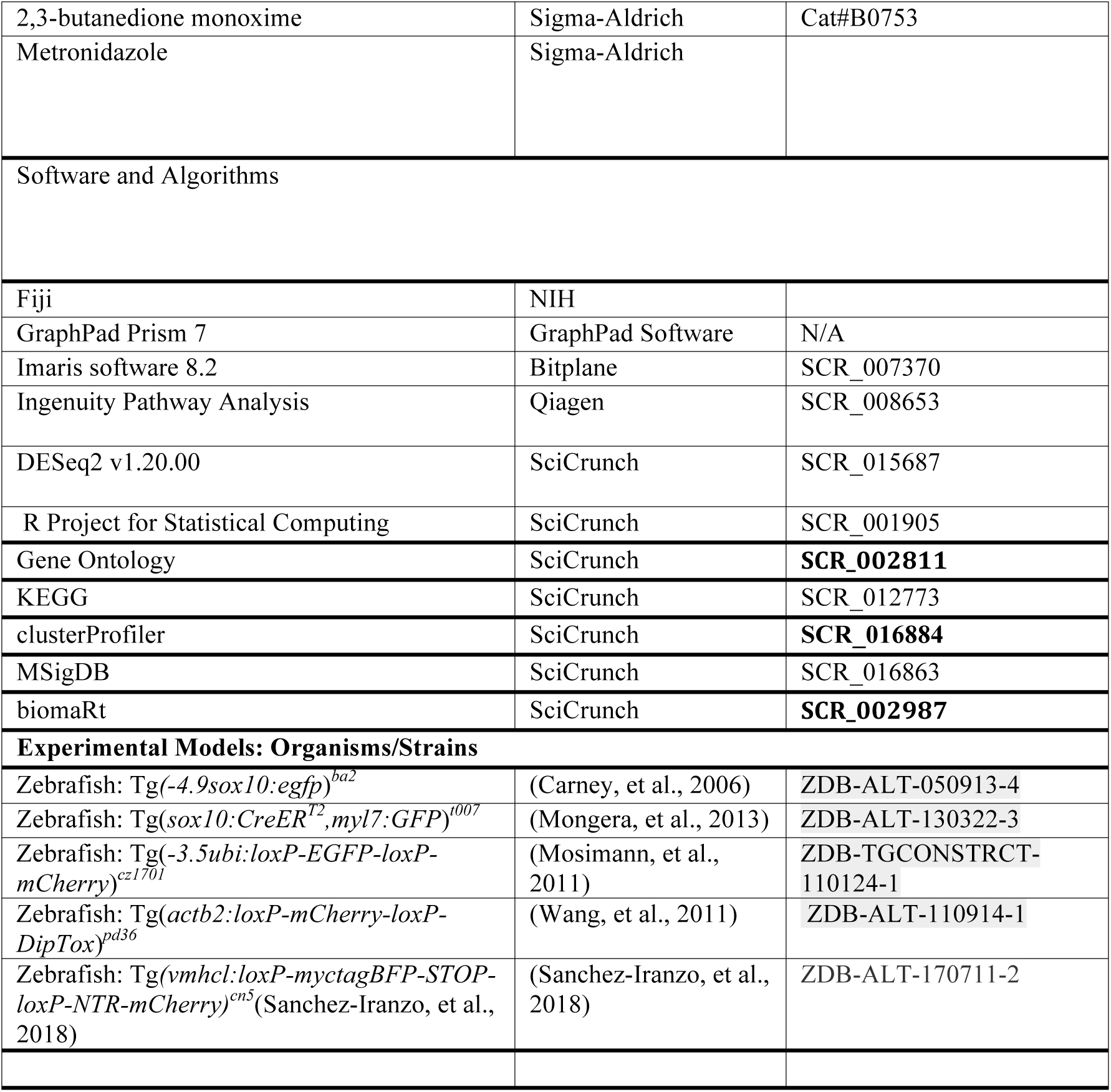

| REAGENT or RESOURCE | SOURCE | IDENTIFIER |
| --- | --- | --- |
| Antibodies |  |  |
| Mouse monoclonal anti MF20 | DSHB | Cat# MF 20,<br>RRID:AB_2147781 |
| Mouse monoclonal anti tropomyosin | DSHB | Cat# CH1,<br>RRID:AB_2205770 |
| Mouse monoclonal anti BrdU | BD Pharmingen | Cat# 347583,<br>RRID:AB_400327 |
| Rat monoclonal anti mCherry | Thermo Fisher Scientific | Cat# M11217,<br>RRID:AB_2536611 |
| Rabbit polyclonal anti-Tyrosine Hydroxylase | Sigma Aldrich | Cat# T8700,<br>RRID:AB_1080430 |
| Rabbit polyclonal anti-RFP | Abcam | Cat# ab34771,<br>RRID:AB_777699 |
| Biotin-SP-conjugated AffiniPure anti-rabbit IgG (H+L) | Jackson Immuno Research Laboratories | Cat# 713-065-003,<br>RRID:AB_2340715 |
| Goat anti-Rabbit IgG (H+L) Secondary Antibody, Alexa Fluor® 568 conjugate | Thermo Fisher Scientific | Cat# S-11226,<br>RRID:AB_2315774 |
| Goat anti-Mouse IgG1 Secondary Antibody, Alexa Fluor® 488 conjugate | Thermo Fisher Scientific | Cat# A-21121;<br>RRID:AB_2535764 |
| Goat anti-Mouse IgG1 Secondary Antibody, Alexa Fluor® 568 conjugate | Thermo Fisher Scientific | Cat# A-21124;<br>RRID:AB_2535766 |
| Goat anti-Mouse IgG1 Secondary Antibody, Alexa Fluor® 647 conjugate | Thermo Fisher Scientific | Cat# A-21240;<br>RRID:AB_2535809 |
| Goat anti-Mouse IgG2b Secondary Antibody, Alexa Fluor® 488 conjugate | Thermo Fisher Scientific | Cat# A-21242,<br>RRID:AB_253581 |
| Chemicals, Peptides, and Recombinant Proteins |  |  |
| 4',6-Diamidino-2-phenylindole dihydrochloride (DAPI) | Thermo Fisher Scientific | Cat# D3571 |
| 5-bromo-2'-deoxyuridine (BrdU) | Sigma Aldrich | Cat# 59-14-3 |
| 4-hydroxytamoxifen | Sigma Aldrich | Cat#T5648 |
| 2,3-butanedione monoxime | Sigma-Aldrich | Cat#B0753 |
| Metronidazole | Sigma-Aldrich |  |
| Software and Algorithms |  |  |
| Fiji | NIH |  |
| GraphPad Prism 7 | GraphPad Software | N/A |
| Imaris software 8.2 | Bitplane | SCR_007370 |
| Ingenuity Pathway Analysis | Qiagen | SCR_008653 |
| DESeq2 v1.20.00 | SciCrunch | SCR_015687 |
| R Project for Statistical Computing | SciCrunch | SCR_001905 |
| Gene Ontology | SciCrunch | **SCR_002811** |
| KEGG | SciCrunch | SCR_012773 |
| clusterProfiler | SciCrunch | **SCR_016884** |
| MSigDB | SciCrunch | SCR_016863 |
| biomaRt | SciCrunch | **SCR_002987** |
| **Experimental Models: Organisms/Strains** |  |  |
| Zebrafish: Tg(-4.9sox10:egfp)<sup>ba2</sup> | (Carney, et al., 2006) | ZDB-ALT-050913-4 |
| Zebrafish: Tg(sox10:CreER<sup>T2</sup>,myl7:GFP)<sup>t007</sup> | (Mongera, et al., 2013) | ZDB-ALT-130322-3 |
| Zebrafish: Tg(-3.5ubi:loxP-EGFP-loxP-mCherry)<sup>cz1701</sup> | (Mosimann, et al., 2011) | ZDB-TGCONSTRCT-110124-1 |
| Zebrafish: Tg(actb2:loxP-mCherry-loxP-DipTox)<sup>pd36</sup> | (Wang, et al., 2011) | ZDB-ALT-110914-1 |
| Zebrafish: Tg(vmhcl:loxP-myctagBFP-STOP-loxP-NTR-mCherry)<sup>cn5</sup> (Sanchez-Iranzo, et al., 2018) | (Sanchez-Iranzo, et al., 2018) | ZDB-ALT-170711-2 |

### Zebrafish husbandry

Experiments were conducted with zebrafish embryos and adults aged 6–18 months, raised at maximal 5 fish/l and maintained under the same environmental conditions: 27.5-28° C, 650-700µs/cm, pH 7.5, the lighting conditions were 14:10 hours (light: dark) and 10% of water exchange a day. Furthermore, all of them were fed three times per day: once-artemia (Ocean Nutrition) and twice-dry food (ZM-000 for larvae stage and Gemma Micron 150 and 300 for juveniles and adults stages, respectively). Experiments were approved by the Community of Madrid “Dirección General de Medio Ambiente” in Spain, the Landesamt für Verbraucherschutz Thüringen, Germany and the “Amt für Landwirtschaft und Natur” from the Canton of Bern, Switzerland. All animal procedures conformed to EU Directive 86/609/EEC and Recommendation 2007/526/EC regarding the protection of animals used for experimental and other scientific purposes, enforced in Spanish law under Real Decreto 1201/2005. Experiments in Switzerland were conducted under the license BE95/15. For longitudinal experiments, the selected animals were grown together with Casper(White, et al., 2008) zebrafish until heart collection at the density as explained above.

### 4-Hydroxytamoxifen administration

4-hydroxytamoxifen (4-OHT; 10 µM; Sigma H7904) was administered at the indicated times and treatments were performed overnight. Prior to administration, the 10 mM stock (dissolved in ethanol) was heated for 10 minutes at 65 °C (Felker, et al., 2016). For genetic labeling in Tg(*sox10:CreER^T2^; ubb:Switch*) embryos, 4-OHT was administered at 10 µM from 12 to 48 hours post-fertilization (hpf). For lineage tracing studies in adult fish, 4-OHT was administered at 10 µM overnight. For genetic ablation using Metronidazole (Mtz; Sigma, M3761), Mtz was diluted in fish water at 10 mM and administered overnight.

### Cryoinjury and analysis of the injured area

Cryoinjury was performed as previously described (González-Rosa and Mercader, 2012). For analysis of regeneration, animals were euthanized at different times post-injury by immersion in 0.16% Tricaine (Sigma, St Louis, MO, USA), and hearts were dissected in media containing 2 U/ml heparin and 0.1 M KCl. For quantification of injured area on adult paraffin heart sections as shown in Fig 1f-g, color deconvolution tool and color threshold tool (Image J Software) were used to segment and measure the injured and uninjured myocardium in µm^2^.

### Metronidazole administration

For genetic ablation using Metronidazole (Mtz; Sigma, M3761), Mtz was diluted in fish water at 10 mM with DMSO at 0.2% and administered overnight.

### BrdU administration

Adult Tg(*sox10:CreER^T2^*;*ubb:Switch*) zebrafish were used to analyze CM proliferation upon cryoinjury. Animals were injected intraperitoneally either at 6 dpi or 29 dpi with 30 µl of 2.5 mg/ml of 5-Bromo-2-deoxyuridine (BrdU, B5002-1G, Sigma). Hearts were collected and processed for analysis at 7 dpi and 30 dpi. To calculate the proliferation index, cryosections were immunostained with anti-BrdU, anti-RFP, and anti-MHC antibodies as described below. At least 3 ventricular sections were imaged for each heart. MHC^+^/mCherry^+^/BrdU^+^ CMs compared to MHC^+^/mCherry^-^/BrdU^+^ CMs were counted manually using Image J software.

### Histological staining

Hearts were fixed in 2 or 4 % paraformaldehyde (PFA) in phosphate-buffered saline (PBS) overnight at 4°C. Samples were then washed in PBS, dehydrated through graded alcohols, washed in Xylol and embedded in paraffin wax. All histological staining were performed on 7 μm paraffin sections cut on a microtome (Leica and Reichert-Jung), mounted on Superfrost slides (Fisher Scientific), and dried overnight at 37°C. Sections were deparaffinized in xylol, rehydrated and washed in distilled water. Connective tissue was stained using Acid Fuchsine Orange G (AFOG) (Gonzalez-Rosa, et al., 2014). Image J software was used to quantify cryoinjured area in uninjured and injured hearts.

### Cardiac imaging by echocardiography

Animals were anaesthetized by immersion for approx. 5 min in a combined solution of 60 mM Tricaine/3 mM Isoflurane dissolved in fish tank water (Gonzalez-Rosa, et al., 2014). Individual fish were placed ventral side up on a custom-made sponge in a Petri dish filled with the anesthetic solution. Two-dimensional (2D) high-resolution real-time *in vivo* images were obtained with the Vevo2100 Imaging System through a RMV708 (22-83 MHz) scanhead (VisualSonics, Toronto, Canada). Imaging and image analysis was performed as described (Gonzalez-Rosa, et al., 2014).

### Immunofluorescence on sections

Heart sections were deparaffinized, rehydrated and washed in distilled water. Epitope retrieval was carried out by boiling in citrate buffer (pH 6.0) for 20 min in a microwave at full power. Sections were permeabilized with Triton X-100 0.5% for 15 min. Non-specific binding sites were saturated by incubation for 1 hour in blocking solution (5% BSA, 5% goat serum, 0.1% Tween-20). Endogenous biotin was blocked with the avidin-biotin blocking kit (Vector, Burlingame, CA, USA). For tyramides amplification, slides were blocked in 3% H_2_0_2_-PBS for 20 minutes. Slides were incubated overnight with the following primary antibodies at 4°C: anti-myosin heavy chain (MF20, DSHB; diluted 1:20), anti-tropomyosin (CH1, DSHB; diluted 1:20), anti-RFP (Abcam, diluted 1:150), anti-BrdU (BD Pharmingen diluted 1:100), anti-mCherry (16D47, Thermo Fisher Scientific) and anti-Tyrosine Hydroxylase (Sigma, diluted 1:150), anti-biotin (Thermo Fisher Scientific, 1:150), HRP (DAKO, 1:250). Antibody signals were detected with biotin- or Alexa (488, 568, 633)-conjugated secondary antibodies (Invitrogen; each diluted 1:250) and streptavidin-Cy3 (Molecular Probes, SA1010) after incubation for 1 hour at room temperature. For tyramides amplification (Merck, TSA Plus Cyanine 3 System, Cat# NEL744001KT), Cy3 was conjugated for 4 minutes. Nuclei were stained with 4’,6-Diamidino-2-phenylindole dihydrochloride (DAPI) (1:1000, Merck) and slides were mounted in DAKO fluorescent mounting medium (DAKO). Images were analyzed and processed using ImageJ.

Cryosections were prepared as above with the following modifications: heart sections were incubated for 30 min in PBS at 37 °C to remove the gelatin. They were washed two more times with PBS at room temperature and then immunofluorescence proceeded as for paraffin sections.

### Whole mount heart imaging and image processing

Adult zebrafish hearts were fixed in 2% PFA overnight. We used CUBIC reagent (Susaki, et al., 2015) for tissue clearing. Hearts were incubated in CUBIC reagent 1 for three days at 37°C, washed in 0.1% PBS/Tween 20 three times for 20 minutes, whole mount immunofluorescence was performed and samples were incubated afterwards in CUBIC reagent 2 for three further days at room temperature. Hearts were mounted on a glass bottom culture dish (MatTek Corporation) for confocal acquisition. Whole heart images were obtained with Zeiss LSM 780, Zeiss LSM 880, and Leica TCS SP8 confocal microscopes with a 10 dry, 20× dry and 40x water dipping lenses. Images were recorded at 512×512, 1024×1024, 2048x 2048 resolution. Tile scan and z-stack of each heart was acquired. The proportion of mCherry^+^/MHC^+^ versus all MHC^+^ CMs was evaluated with Imaris software 8.2 (BITPLANE). A distance transformation algorithm (Imaris software 8.2) was used to study the distance of mCherry^+^/MHC^+^ CMs to the injured or apex area.

### Imaging of larvae *in vivo*

Double transgenic larvae Tg(*sox10:CreER^T2^*;*ubb:Switch*) were transferred to E3 medium containing 0.2 mg/ml tricaine and 0.0033 % PTU and immobilize using 0.7% agarose (Bio-Rad Low Melting agarose, #Cat 161-3111) in a glass bottom microwell-dish (MatTek Corporation). Zebrafish hearts were scanned using bidirectionally acquisition with SP5 confocal microscope (Leica SP5) using a 20x glycerol lens. Larvae were carefully removed from the agarose embedding and were grown to adults in fish tanks together with Casper fish. Adult zebrafish hearts were collected, fixed in 2% PFA overnight and scanned with LSM 700 Confocal microscope using 20x dry lens (Zeiss). 3D reconstruction and analysis were done using Imaris Software 8.2.

### RNAscope 2.5HD Detection Reagent (RED) - Immunofluorescence method

All the hearts were fixed at room temperature (RT) for 24h in 10% Neutral Formalin Buffer (NFB). After the fixation, samples were washed 3 times for 10 minutes in 1x PBS. Dehydration process was performed using a standard ethanol series (10 minutes each), followed by two xylene washes (5 minutes each) and embedding of tissues was carry out.

Paraffin blocks were cut with microtome (Microm) at 6µm thickness per section, collected in the water bath with SuperFrost slides and baked slides in a dry oven for one hour at 60° C. After that, deparaffination process was done: 2 times for 5 minutes in xylene, 2 times for 2 minutes in 100% ethanol and dried slides in a dry oven for 5 minutes at 60°C. Then, permeabilization with hydrogen peroxide (ACD#322381) was accomplished for 10 minutes at RT and washed 2 times for 1 minute in distilled water.

Target retrieval (ACD, #322000) was performed for 15 minutes at 100° C, next washed for 15 seconds in distilled water and 100% ethanol for 3 minutes, then dry slides for 5 minutes at 60° C and a hydrophobic barrier was created for every section with PAP Pen (Vector, #H-4000). After that, protease Plus (ACD#322381) for 5 minutes at 40° C and washed slides in distilled water two times.

Probes were designed for the experiment (Dr-Sox10 and negative control probe-DapB) and hybridization was accomplished with the incubation of the probes for 2 hours at 40° C and 2 times for 2 minutes in wash buffer (ACD, #310091).

RNAscope 2.5 HD Detection Reagent-RED (ACD, #322360) were used next: AMP1 (30min at 40° C)-wash buffer-AMP2 (15min at 40° C)-wash buffer-AMP3 (30min at 40° C)-wash buffer-AMP4 (15min at 40° C) - wash buffer-AMP5 (30min at 40° C)-wash buffer-AMP6 (15min at 40° C) - wash buffer-RED working solution (10min at RT)-wash in distilled water (2 times for 5 minutes).

### Statistical analysis

The specific test used in each comparison is indicated in the main text or figure legend. Normal distribution was tested to decide if a parametric or non-parametric test needed to be applied.

### Disaggregation of zebrafish hearts, cardiomyocytes sorting and RNA-Seq library production

Uninjured and injured recombined adult zebrafish *sox10:CreER^T2^*;*vmhcl:loxP-tagBFP-loxP-mCherry-NTR* hearts were collected 12 days after the last 4-OHT pulse or 7 dpi respectively, and processed according to previous protocols(Sanchez-Iranzo, et al., 2018; Tessadori, et al., 2012). Atrium and *bulbus arteriosus* were removed to obtain only the ventricle. Three adult zebrafish ventricles were pooled for each sample and pools of 20 CMs each sorted.

mCherry^+^, mCherry;BFP^+^ and BFP^-^ CMs were sorted in 0.2 ml tubes in lysis buffer using Synergy 4L Cell Sorter and immediately frozen at −80°C. Smart-Seq2 RNA library preparation was performed according to previous protocols(Picelli, et al., 2014). An Agilent Bioanalyzer was used to measure quality of library preparation. Library concentration was measured using the Qubit fluorometer (ThermoFisher Scientific). Final libraries concentration was 10 nM. Libraries were sequenced using Illumina NextSeq 500.

### Bioinformatics analysis

BCL files were converted to FastQ files, using bcl2fastq2 (v2.20.0.422 – Illumina). Reads were mapped to the reference genome (Ensembl build 11, release 94) using Hisat2, version 2.1.0(Kim, et al., 2015) and counting was performed using featureCounts, version 1.6.0(Liao, et al., 2014) Multiple quality control features were measured and observed using both FastQC, version 0.11.5(Andrews, 2010) and RseQC, version 2.6.4 (Wang, et al., 2012) From 44 sequenced samples 18 were discarded as they did not pass the quality control. For bioinformatics analysis, we compared mCherry^+^ (mCherry^+^, mCherry^+^BFP^+^) with mCherry^-^(BFP^+^) pools. The comparison was performed for samples extracted from uninjured and 7 dpi hearts. Downstream analysis was performed in R, version 3.5.1 (R Core Team (2018). R: A language and environment for statistical computing. R Foundation for Statistical Computing, Vienna, Austria. URL https://www.R-project.org/.).

Counts were normalized and differential expression between design groups was tested using package DESeq2 v.1.20.00 with no log2 fold change shrinkage (default betaPrior option for the latest versions of the tool). Principal component analysis (PCA) plots, volcano plots and heatmaps were generated using the ggplot2 package, version 3.0.0 (H. Wickham. ggplot2: Elegant Graphics for Data Analysis. Springer-Verlag New York, 2016.).

Further analyses were performed with DESeq2 results. For the enrichment we selected all the Gene Stable IDs and translated to *Mus musculus* Gene Stable IDs and obtained the ENTREZIDs and SYMBOLs using biomaRt package (Durinck, et al., 2005). With the genes translated, the top differentially expressed genes (DEG) that passed FDR < 0.05 for Gene Ontology (GO)(Ashburner, et al., 2000) over representation analysis (ORA) using clusterProfiler package (Yu, et al., 2012).

Afterwards, we performed deeper analysis for overall gene expression with gene set enrichment analysis (GSEA). The differential gene expression results from DESeq2, were sorted by Log2FoldChange value, providing a ranking and a DGE direction within the comparison. Kyoto Encyclopedia of Genes and Genomes (KEGG)(Kanehisa and Goto, 2000) and Molecular Signature Database (MsigDB)(Liberzon, et al., 2015) using the Hallmarks collection were used for biological insight. For the GSEA analyses clusterProfiler and FGSEA packages (Sergushichev, 2016) were used with KEGG and MsigDB Hallmarks gene set respectively. For data representation only those with adjusted p-value < 0.05 were considered significant of the results obtained.

Ingenuity pathway analysis core analysis (IPA, QIAGEN Inc., https://www.qiagenbioinformatics.com/products/ingenuity-pathway-analysis) was used to identify canonical pathways, functions and diseases related to our differentially expressed genes in uninjured and injured conditions.

## Author contributions

M.S. performed the experiments, prepared Figures and contributed to manuscript writing, I.M. performed experiments in Fig. 1, S1 and supervised M. P.-L. M.P.-L. contributed to data shown in Fig. 1,S1 M.P.-L and I.M. made the initial observation of a contribution of sox10-dervied cells to regeneration. M.G.-C. helped with experiments in Fig. 4 and S2-S6. X.L. helped with experiments shown in Fig. 3, 5 and S2 and was responsible for fish line maintenance. G. G.-M. contributed to Figure S6. M.B. helped with bioinformatics analysis in Fig.5 and S3-5. D.P. and V.B. contributed to RNA-seq. D.F. and R.B. contributed to bioinformatics analysis. N.M. designed and interpreted the experiments, wrote the manuscript and secured funding.

## Acknowledgements

We are grateful to Tilly Mommersteeg for sharing unpublished results, Lorena Flores and Ana Vanesa Alonso for help with echocardiography, Jonathan Landry for bioinformatics advice and Ronja Baal for fish husbandry. We thank the Animal facility, Histology, Cellomic and Microscopy Units from CNIC, the MIC-Bern Unit and Cellomics Unit at the University of Bern for support with experiments and Ken Poss, Robert Kelsh and Alessandro Monguera for sharing fish lines. We thank JM González-Rosa and Fernando Rodríguez-Pascual for fruitful discussion. This work has been funded by the Spanish Ministry of Economy and Competitiveness through grant BFU2014–56970–P to N.M. and SAF2012-34916 y SAF2015-65679R to F.R-P., the Swiss National Science Foundation grant 31003A_159721, the ERC starting grant 337703–zebra–Heart, Grupos de investigación de la Comunidad de Madrid en Biomedicina, (FIBROTEAM S2010/BMD-2321), and co-funding by Fondo Europeo de Desarrollo Regional (FEDER). I.J.M. was supported by PIEF-GA-2012-330728. D.F. was supported by the European Commission Marie Sklodowska-Curie Innovative Training Network: BtRAIN – European Brain Barriers Training Network (H2020-MSCA-ITN-2015, n°675619)”. The CNIC is supported by the Spanish Ministry of Economy and Competitiveness (MINECO) and the Pro-CNIC Foundation, and is a Severo Ochoa Center of Excellence (MINECO award SEV-2011-0052 and SEV-2015-0505).

---

**Figure S1.**
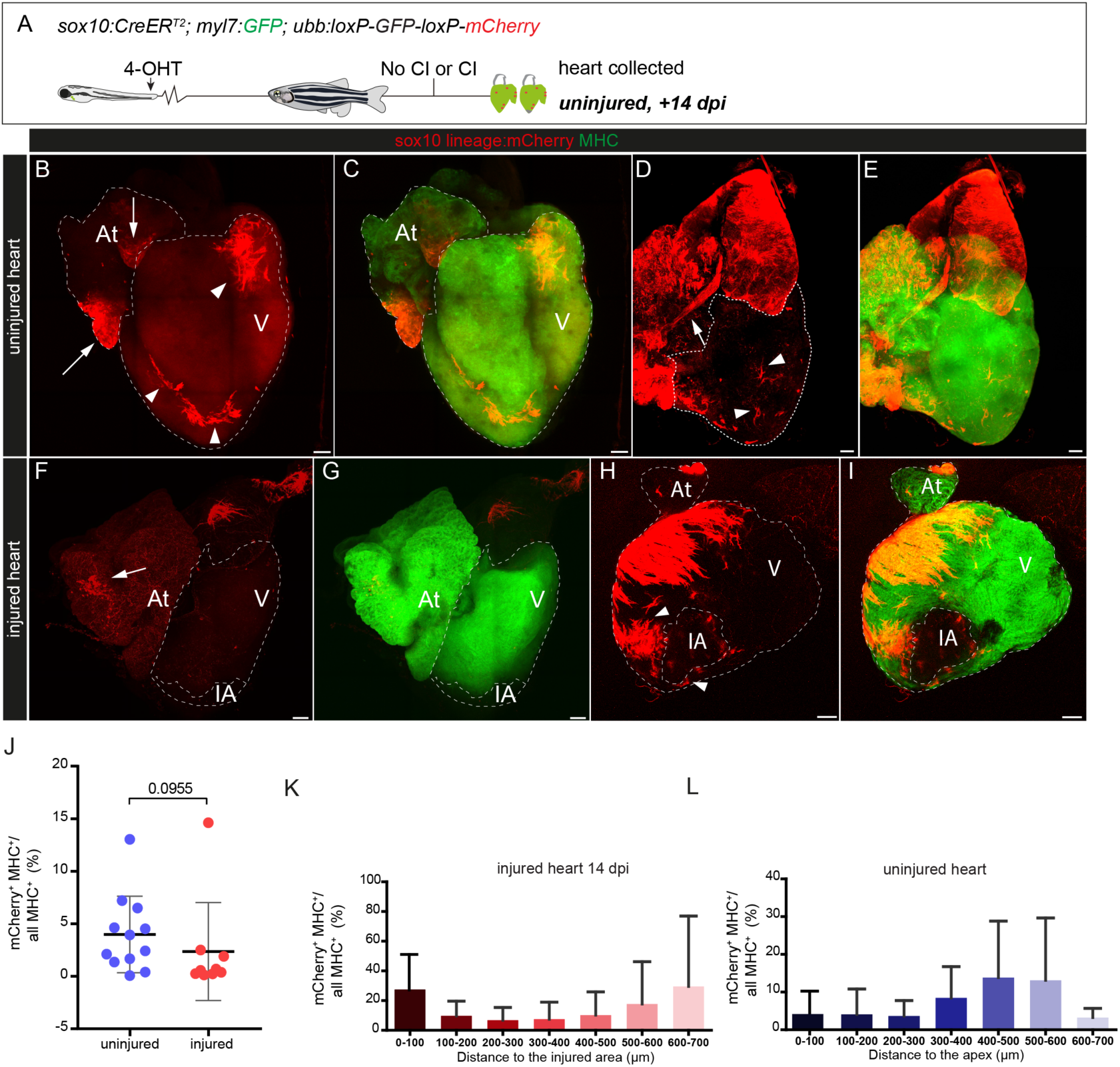
Embryonic *sox10*-derived CMs do not expand after cryoinjury. Related to Fig. 1. (A) 4-OHT was administered to *sox10:CreER^T2^;ubb:loxP-GFP-loxP-mCherry* zebrafish during embryogenesis (12–48 hours post-fertilization) and adult uninjured hearts or hearts at 14 dpi were collected and imaged. (B-I) Maximal intensity projection of a confocal z-stack through a heart. *sox10*-derived cells are mCherry^+^ (red). The whole myocardium is MHC^+^ (green). mCherry^+^ cells are present in the ventricle (arrowheads) and atrium (arrows). (J) Relative volume of embryonic *sox10*-derived CMs (mCherry^+^/ MHC^+^) compared to all CMs (MHC^+^) in uninjured (n= 12) and injured hearts at 14 dpi (n=9). Shown are values from individual hearts as well as means ± s.d.; p=0.0955 (two-tailed non-parametric t-test). (K,L) Quantification of the distribution of mCherry^+^ CMs in whole mount immunostained hearts. The ventricle was digitally sectioned in increments of 100 µm starting from the injury site or apex (0). Shown is the relative volume from mCherry^+^/MHC^+^ cells versus all MHC^+^ cells within different heart segments. The amount of mCherry^+^/MHC^+^ does not differ between segments (one-way ANOVA). n= 5 hearts for 14 dpi and n=5 for uninjured condition. Atrium, ventricle and injury area are outlined by dotted lines. 4-OHT, 4-hydroxytamoxifen; At, atrium; dpi, days post injury; IA, injured area; CI, cryoinjury; CMs, cardiomyocytes; V, ventricle. Scale bars: 100 µm.

**Figure S2.**
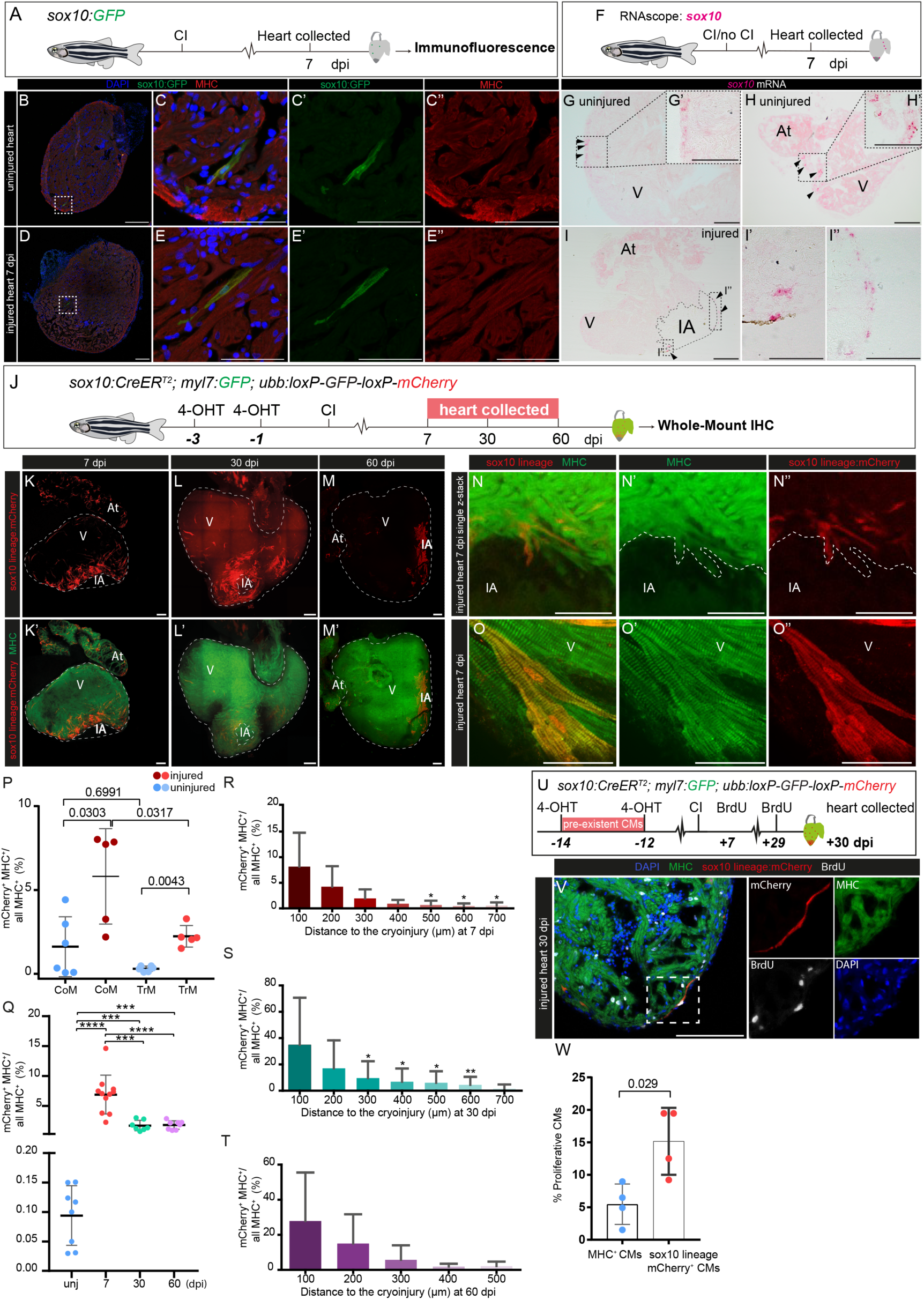
*sox10*-derived CMs participates in the heart regeneration. Related to Figure 3. (A-E’’) Detection of a small subset of *sox10*:GFP^+^ CMs in the adult zebrafish heart. (A) Immunostaining against GFP (green) Myosin Heavy chain (MHC, red) and nuclear counterstain with DAPI (blue) was performed on sagittal sections of *sox10:eGFP* adult zebrafish hearts. (B-C’’) Whole heart section and zoomed view of an uninjured heart. (D-E’’) Heart at 7 dpi. For both heart sections a GFP^+^ ventricular CM is shown. GFP, green fluorescence protein; The cardiac ventricles were collected on 2 slides. Number of GFP+ cells were counted on 1 slide per heart. The number of *sox10*:GFP^+^ CMs detected on sections ranged between 0 to 14 for both conditions. (F-I’’) RNAScope for *sox10* mRNA on sections of uninjured hearts and hearts at 7dpi. Arrowheads, *sox10* positive cells. (J-T) Analysis of the contribution of *sox10*-derived cells present in the uninjured heart to heart regeneration using *sox10:CreER^T2^;ubb:loxP-GFP-loxP-mCherry*. (J) Two 4-OHT treatments were performed on alternating days to adult *sox10:CreER^T2^;ubb:loxP-GFP-loxP-mCherry* zebrafish before heart cryoinjury. Hearts were collected at 7 (n=11), 30 (n=7) and 60 (n=9) dpi. (K-O) Whole-mount views of an injured hearts revealing *sox10*-derived mCherry^+^ cells in red and the myocardium in green (MHC^+^ expression). (K,K’) Heart at 7 dpi. *sox10*-derived cells surround the injury area. (L,L’) Heart at 30 dpi. *sox10*-derived CMs are present in the injured area and border zone. (M,M’) Heart at 60 dpi. *sox10*-derived cells are found within the regenerated myocardium. (N-N’’) Zoomed view of a 8 µm z-stack from a heart at 7 dpi. (O-O’’) Magnifications showing *sox10*-derived CMs at 7 dpi. (P) Contribution of *sox10*-derived CMs to the trabecular or compact myocardium. Graph showing the percentage of mCherry^+^/MHC^+^ versus all MHC^+^ cells in the compact (comp) or trabecular (trab) myocardium in injured and uninjured hearts. Dots represent measurements from individual hearts, Dots represent measurements from individual hearts, shown are also means ± s.d (p=0.0303 and 0.0043 two-tailed non-parametric t-test). For this experiment, recombination was performed at −12 and −14 days before cryoinjury. (Q) Percentage of the volume from mCherry^+^/MHC^+^ cells relative to all MHC^+^ cells in uninjured and injured hearts. The percentage of *sox10*-derived CMs volume significantly expands upon injury. ***p<0.001, ****p<0.0001 (according to one-way ANOVA followed by Tukey’s honest significant difference test). (R-T) Graph showing the percentage of the volume from mCherry^+^/MHC^+^ cells relative to all MHC^+^ cells within different heart segments at 7, 30 and 60 dpi. Data are means ± s.d.; *p<0.05; **p<0.01; ***p<0.001 (according to one-way ANOVA followed by Tukey’s honest significant difference test). (U-W) Analysis of BrdU incorporation to determine *sox10*-derived CM proliferation in *sox10:CreER^T2^;ubb:loxP-GFP-loxP-mCherry* zebrafish. 4-OHT was added at −12 and −14 days before cryonjury. BrdU was added at 6 and 29 dpi. Hearts were collected at 30 dpi. (V) Immunofluorescence staining on heart at 30 dpi. MHC, green; mCherry, red; BrdU, white; nuclei are counterstained with DAPI, blue. Shown are merged imaged and single channels and zoomed region is highlighted with dotted lines. (W) Quantification of BrdU^+^ CMs at 30 dpi. *sox10*-derived CMs have a higher proliferative capacity than other CMs in the zebrafish heart. Data means ± s.d (p=0.029; two-tailed non-parametric t-test). 4-OHT, 4-hydroxytamoxifen; At, atrium; DAPI, 4’,6-diamidino-2-phenylindole; dpi, days post injury; IA, injured area; CI, cryoinjury; CMs, cardiomyocytes; MHC, myosin heavy chain; V, ventricle. Scale bars: 100 µm; C-C’’,E-E’’ and O-O’’’ 50 µm.

**Figure S3.**
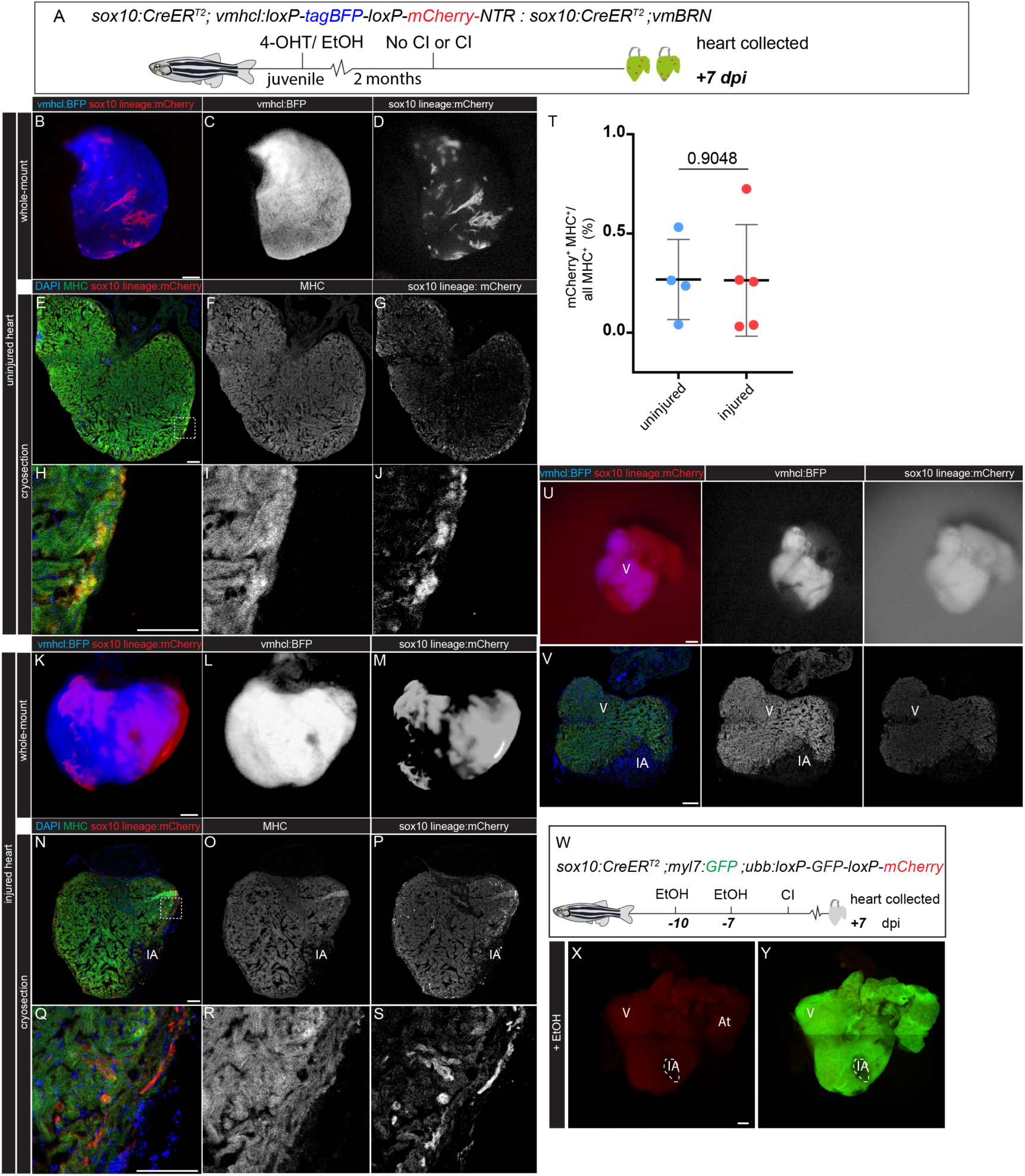
Recombination in juvenile *sox10*:CreERT2 zebrafish and control of 4-OHT recombination induction. Related to Figure 3. (A) 4-OHT was administered to *sox10:CreER^T2^;ubb:loxP-GFP-loxP-mCherry* zebrafish during juvenile stage (4 weeks post-fertilization) and adult uninjured hearts or hearts at 7 dpi were collected and imaged. (B-D) Fluorescence stereomicroscope acquisition of a dissected heart showing *sox10*-derived cells in red (mCherry^+^) and ventricular CMs in blue (*vmhcl*:BFP^+^) in uninjured conditions. (E-J) Cryosection of an uninjured hearts. *sox10*-derived CMs are detected near the subepicardial regions of the myocardium. (H-J) Zoomed views of boxed regions in E. (I) Showing MHC^+^ cells and (I) *sox10*-derived CMs (mCherry^+^, J). (K-M) Fluorescence stereomicroscope acquisition of a dissected heart showing *sox10*-derived cells in red (mCherry^+^) and ventricular CMs in blue (*vmhcl*:BFP^+^) in injured conditions. (N-S) Cryosection of an injured heart at 7 dpi.*sox10*-derived CMs are detected near the subepicardial regions and close to the injured area. (Q-S) Zoomed views of boxed regions in N. Shown are merged and single channels of MHC and mCherry signal. (T) Percentage of juvenile *sox10*-derived CMs area (mCherry^+^/ MHC^+^) compared to area of all myocardial cells (MHC^+^) in uninjured (n= 4) and injured hearts at 7 dpi (n=5). Data means ± s.d.; p=0.9048 (two-tailed non-parametric t-test). (U-V) *sox10:CreER^T2^;ubb:loxP-GFP-loxP-mCherry* zebrafish as shown in K-M but without 4-OHT treatment. Note the lack of mCherry^+^ cells in whole mount views (upper row) and on sections (lower row) (n=2). (W-Y) EtOH was administered to *sox10:CreER^T2^;ubb:loxP-GFP-loxP-mCherry* zebrafish 10 and 7 days before injury and hearts at 7 dpi were collected and imaged (n=3). mCherry^+^ cells could not be detected. 4-OHT, 4-hydroxytamoxifen; At, atrium; CI, cryoinjury; CMs, cardiomyocytes; dpi, days post-injury; EtOH, ethanol; IA, injured area; V, ventricle. Scale bar: 50 µm.

**Figure S4.**
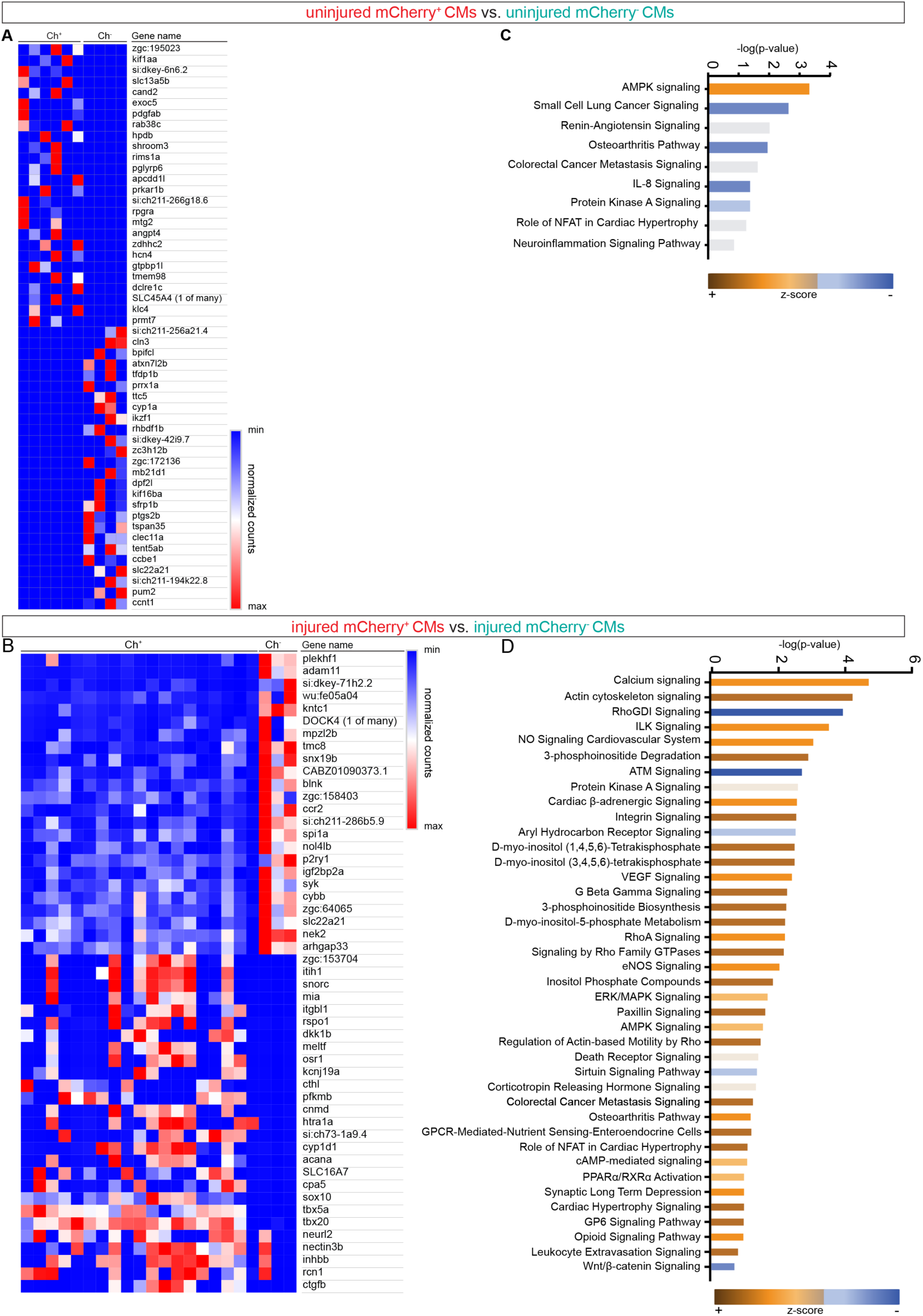
Heatmaps and canonical pathway analysis of *sox10*-derived CMs *vs* rest of CM in injured and uninjured hearts. Related to Figure 4. (A) Heatmap representing the 50 most up or downregulated genes when comparing mCherry^+^ with mCherry^-^ samples of uninjured hearts. Genes are as represented in the volcano plot of Fig. 4. (B) Heatmap representing the 50 most up or downregulated genes when comparing mCherry^+^ with mCherry^-^ samples from injured hearts. Genes are as represented in the volcano plot of Fig. 4. (C) All canonical pathways activated or inhibited according to DEG list when comparing mCherry^+^ with mCherry^-^ CMs from uninjured hearts (Ingenuity pathway analysis (IPA). (D) Top 40 canonical pathways predicted to be activated or de-activated according to DEG between mCherry^+^ and mCherry^-^ CMs at 7 dpi according to IPA. Ch, mCherry; CMs, Cardiomyocytes; dpi, days post injury.

**Figure S5.**
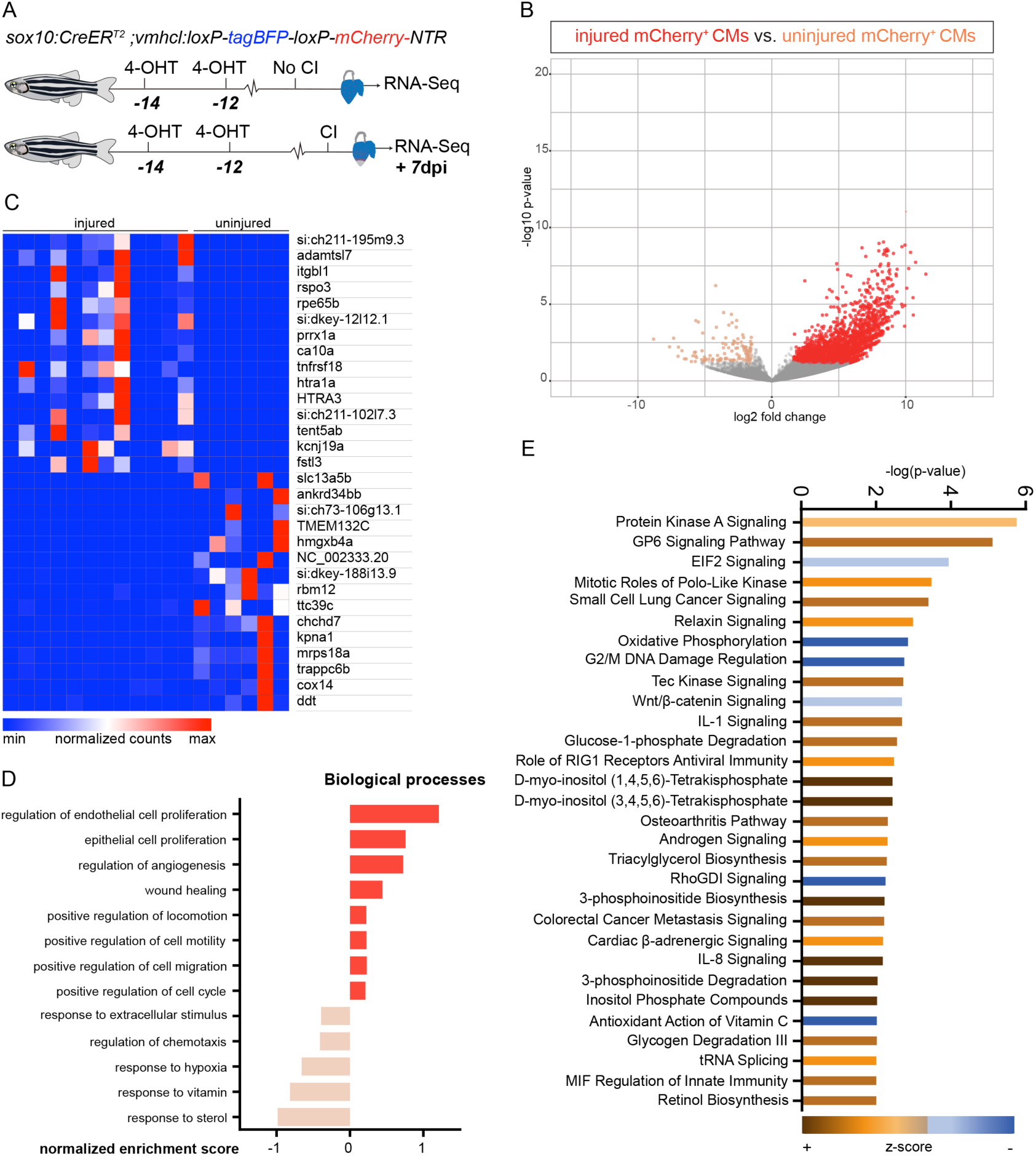
Changes in gene expression of mCherry^+^ (*sox10*-derived) CMs in response to injury. Related to Figure 4. (A) Uninjured zebrafish hearts and hearts at 7dpi from the *sox10:CreER^T2^*;*vmhcl:loxP-tagBFP-loxP-mCherry-NTR* line were collected 12 days after 4-OHT treatment. mCherry^+^ and mCherry^-^ cells were sorted and RNA-seq performed. (B) Volcano plot representing differentially expressed genes (DEG) of Cherry^+^ cells from uninjured and injured hearts. DEG uninjured heart, orange dots. DEG injured heart, red dots. DEG defined as adj p-value ≤0.05 and a log2 fold change (LFC) ≥ 2. Grey dots, genes with an FDR ≥0.05 and LFC ≤ ± 2. A higher number of DEG in mCherry^+^ CMs was observed at 7 dpi. (C) Heatmap representing the 16 most up and 16 most downregulated genes when comparing injured and uninjured mCherry^+^ samples (D) Biological processes of DEG present in Cherry^+^ CMs from uninjured (orange bars) and injured hearts (red bars). Representative biological processes were plotted and ordered according to the normalized enrichment score. Z-score represents whether a specific function is increased or decreased according to the DEG. (E) Most significant canonical pathways activated or inhibited when comparing mCherry^+^ CMs from injured and uninjured hearts using Ingenuity Pathway analysis (IPA). (E). Ch, mCherry; CMs, Cardiomyocytes; dpi, days post injury; GO; Gene Ontology.

**Figure S6.**
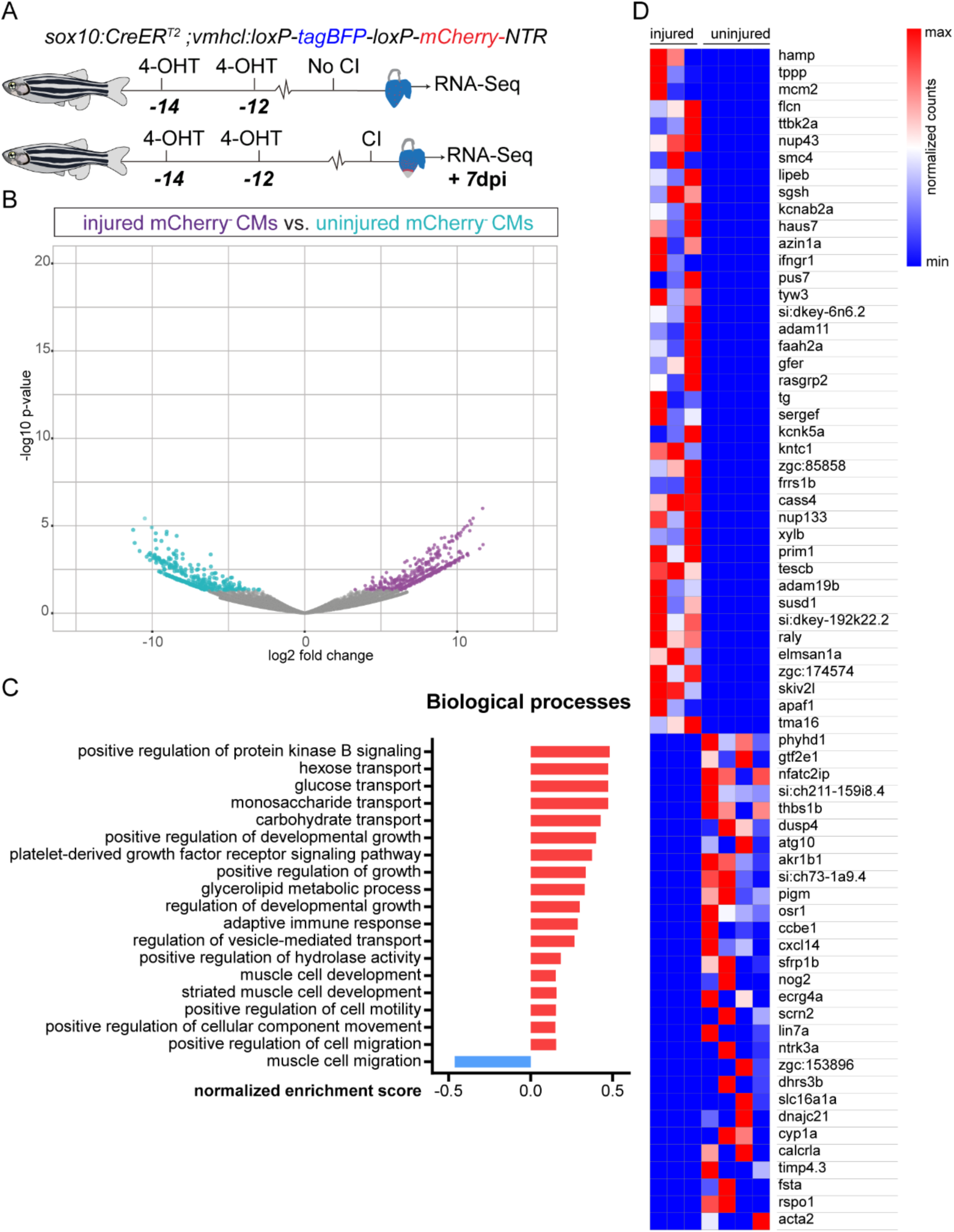
Changes in gene expression of mCherry^-^ CMs (not *sox10*-derived) in response to injury. Related to Figure 4. (A) Uninjured hearts and hearts at 7dpi from the *sox10:CreER^T2^*;*vmhcl:loxP-tagBFP-loxP-mCherry-NTR* line were collected 12 days after 4-OHT treatment. mCHerry^+^ and mCherry^-^ cells were sorted and RNA-seq performed. (B) Volcano plot representing differentially expressed genes (DEG) in Cherry^-^ CMs from uninjured and injured.hearts. Orange dots, DEG win uninjured Cherry^-^ CMs; red dots, injured Cherry^-^ CMs. DEG: adj p-value≤0.05 and a log2 fold change (LFC) ≥ 2. Grey dots, genes with an adj p-value ≥0.05 and LFC ≤ ± 2. Similar number of DEG in the comparison of these two populations in uninjured and injured hearts. (C) GO Biological processes upregulated in injured Cherry^-^ (red bars) and uninjured Cherry^-^ CMs (blue bars). Representative biological processes were plotted and ordered according to the normalized enrichment score. Z-score represents whether a specific function is increased or decreased according to the DEG. (D) Heatmap representing the 43 most upregulated and 29 most downregulated genes when comparing injured and uninjured mCherry^+^ samples. Ch, mCherry; CMs, Cardiomyocytes; dpi, days post injury; GO; Gene Ontology.

**Figure S7.**
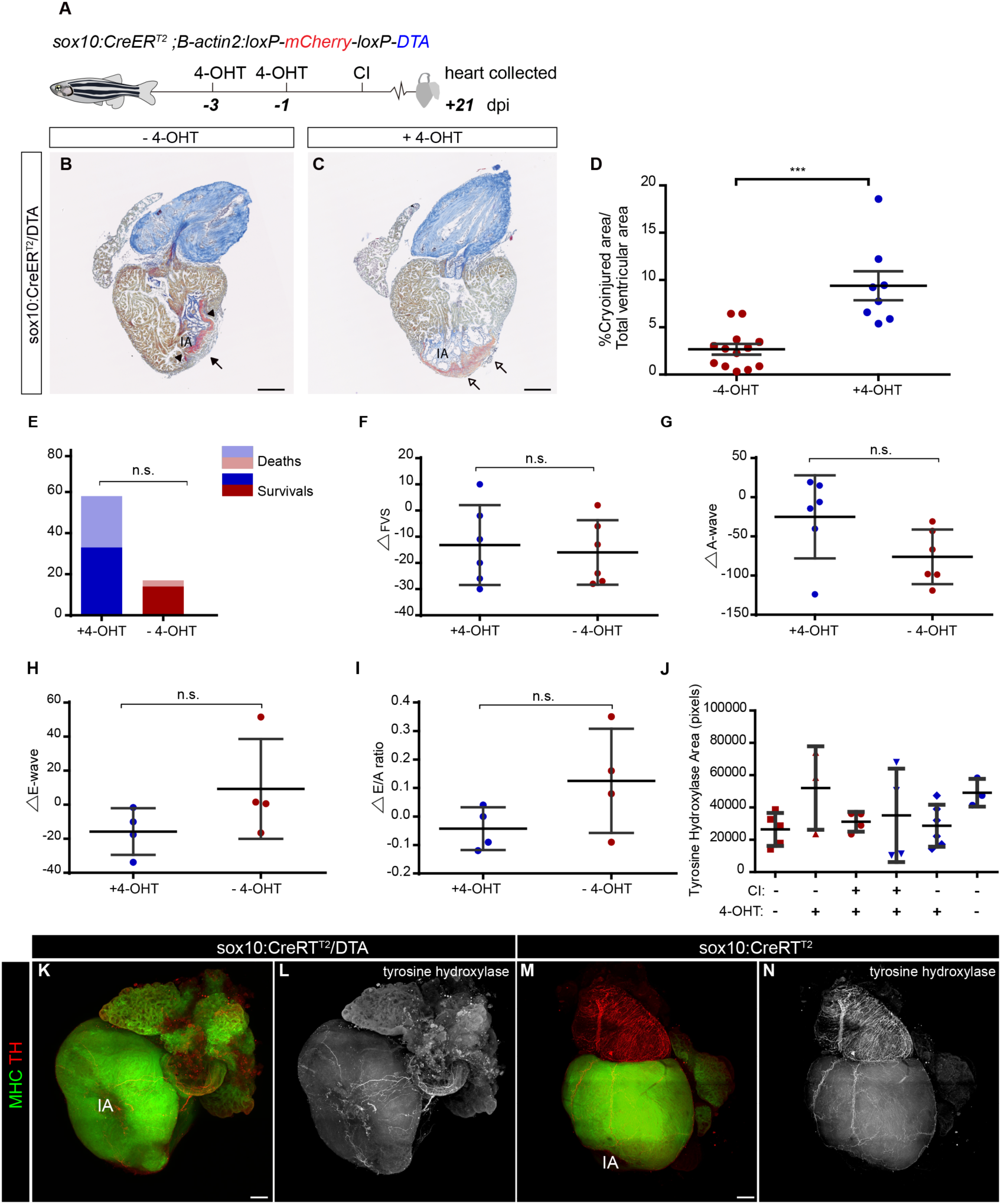
Ablation of adult *sox10*-derived cells does not affect cardiac homeostasis but impairs cardiac regeneration. Related to Figure 5. (A) At day −1 and −3 prior to cryoinjury, *sox10:CreER^T2^*;*bactin*:*loxP-mCherry-loxP-DTA* zebrafish (abbreviated as *sox10:CreER^T2^;DTA*) were either treated with 4-OHT (n=8) or with adjuvants (control group; n=13). Cryoinjured hearts were collected at 21 days post-injury (dpi). (B,C) AFOG histological staining on sagittal heart sections. (B) Control heart revealing collagen deposition close to the injury area (arrowheads) and regenerated myocardial layer (arrows). (C) 4-OHT-treated hearts revealing a larger amount of collagen deposition close to the IA and absence of myocardial regeneration (empty arrows). (D) Quantification of the IA in control hearts and hearts from *sox10*-derived cell depleted animals. Data are means ± s.d.; p= 0.0006 (two-tailed non-parametric t-test). (E) Viability of *sox10:CreER^T2^;DTA* animals with and without 4-OHT administration. Data are means ± s.d.; p= 0.0862 (Fisher’s exact test). (F-I) Echocardiographic measurements. Cardiac function was assessed in 4-OHT-treated animals and untreated controls. Data are means ± s.d.; p values were obtained using a non-parametric t-test. (J-N) Cardiac innervation upon *sox10*-derived cell ablation. (J) Quantification of TH signal. No difference in cardiac innervation was observed in hearts from *sox10:CreER^T2^;DTA* (n=5) and *sox10:CreER^T2^* (n=4) zebrafish according to one-way ANOVA. (K-N) Whole-mount immunofluorescence using anti-tyrosine hydroxylase (TH) and anti-MHC staining on 14 dpi hearts from 4-OHT-treated *sox10:CreER^T2^;DTA*. Shown is a z-Stack comprising 35 z-planes (285 µm) for sox10:CreER^T2^;DTA a z-stack comprising 32 z-planes (260 µm) for the *sox10:CreER^T2^* heart. 4-OHT, 4-Hydroxytamoxifen; At, atrium; dpi, days post-injury; FVS, fractional volume shortening; IA, injured area; MHC, Myosin Heavy Chain; TH, tyrosine hydroxylase. V, ventricle. Scale bars: 100 µm (B and C); 200 µm (K-N).

